# TDP-43 controls RNA structure through high affinity lattice interactions

**DOI:** 10.1101/2025.11.26.690829

**Authors:** Rahul Vivek, Takuma Kume, Saeed Roschdi, M. Thomas Record, Aaron A. Hoskins, Samuel E. Butcher

## Abstract

TDP-43 is an RNA binding protein implicated in neurodegenerative disease. TDP-43 binds to GU dinucleotide repeats, which are highly abundant sequences in human RNA. Here we show TDP- 43 has one of the highest affinities and specificities measured for an RNA binding protein. Binding prevents formation of the pUG fold, an intramolecular quadruplex, and conversely pUG fold formation prevents TDP-43 binding. A rapid on-rate allows TDP-43 to capture single stranded RNA prior to folding. The protein recognizes the RNA as a 1D lattice, in which overlapping binding sites produce efficient initial binding events that interfere with subsequent interactions. This effect is partially overcome by RNA facilitated protein-protein interactions, which serve to increase the on-rate of a second TDP-43 molecule. In conjunction with all atom models, these data reveal how TDP-43 recognizes RNA repeat sequences and identify an interplay between RNA folding and protein recognition that may be relevant to human disease.

## Introduction

TDP-43 is a key RNA-binding protein and its loss of function is associated with the severe neurodegenerative diseases amyotrophic lateral sclerosis (ALS) (1-3), frontotemporal lobar degeneration (FTLD) (1,4), and Alzheimer’s disease (5,6). Multiple crosslinking and immunoprecipitation studies have shown that TDP-43 preferentially binds to UG repeats, also known as poly(UG) or “pUG” RNA sequences in cells (7-10). These pUG RNA motifs can be up to 50 dinucleotide repeats in length (8,10). The interaction of TDP-43 with pUG RNA is crucial for its functions in regulating RNA metabolism in various processes such as alternative splicing (11-13), cryptic exon repression (14-19), condensate formation (20,21), alternative poly-adenylation (22,23), and nuclear retention (24-26). How TDP-43 achieves its broad range of functions is not yet clear but likely involves recruitment of other proteins to RNA (27,28).

When fractionated from cultured human cells, TDP-43 is 96% monomer and 4% dimer (29). Dimerization is promoted by the N-terminal domain (NTD), which self-associates through weak “head to tail” electrostatic interactions with a *K*_D_ of ∼2 µM in the absence of RNA (30-32). At high micromolar protein concentrations, the NTD promotes protein oligomerization *in vitro* (30,33,34). The RNA binding domain (RBD) of TDP-43 consists of 2 RNA recognition motifs that bind to single-stranded RNA, and the NMR structure of the isolated RBD bound to a non-pUG RNA has been described (35). TDP-43 also has a C-terminal domain (CTD) that is predicted to be intrinsically disordered and can form aggregates of insoluble fibers that are a hallmark of ALS/FTLD and are associated with loss of nuclear function (36-39).

Humans have ∼20,000 pUG sequences with 12 or more repeats, which are predominantly found in introns near splice sites (40). We have shown that pUG RNAs with 12 or more repeats fold into a stable left-handed, intramolecular RNA quadruplex (G4) structure, the pUG fold (40-42). This fold contains three G-quartets and a U-quartet coordinated to 3 potassium ions. The pUG fold forms *in vivo* in *C. elegans*, where it promotes gene silencing by recruiting RNA dependent RNA polymerase to RNAs for the synthesis of siRNAs (40,41,43). Previous studies have characterized TDP-43 binding to pUG oligonucleotides *in vitro* (21,35,44-49), but have used either short oligonucleotides that cannot adopt pUG folds, or non-physiological ionic conditions (e.g., Na^+^ buffers without K^+^), that do not support pUG fold formation. TDP-43 can also bind to G4s (50,51), although this binding activity has been attributed to an aggregated state of the protein (52). The binding mechanism of soluble TDP-43 to longer, commonly occurring pUG sequences in physiologically relevant buffers has not been established.

Here we show that soluble TDP-43 lacking the CTD binds tightly to single-stranded pUG RNA, but not to pUG RNAs folded into quadruplexes (pUG folds). A diffusion-limited on-rate enables TDP-43 to capture single stranded pUG RNA prior to pUG folding. The binding site size is 12 nucleotides (nts) and 2 copies of TDP-43 bind non-cooperatively to the commonly occurring 24-nt RNA sequence (GU)_12_. The first binding event is extraordinarily tight with a *K*_D_ ≤ 5 pM. While binding is non-cooperative, the binding isotherm appears anticooperative (i.e., negative cooperativity) because of overlapping binding sites, in which most initial binding events inhibit the binding of a second molecule. We illustrate how this behavior can be quantitatively understood as a 1D lattice interaction. Binding is highly specific and affinity drops by 5 orders of magnitude when 2 dinucleotides are substituted with adenosines. This specificity can be explained by all-atom models derived from AlphaFold. A second TDP-43 binding event is facilitated by RNA-templated protein-protein interactions involving the NTD, which significantly increase the on-rate of a second molecule. Despite these cooperative protein-protein interactions, binding is non-cooperative because of the 1D lattice. These data elucidate a complex TDP-43 pUG RNA binding mechanism and reveal a previously unrecognized role for TDP-43 in preventing intramolecular RNA folding.

## Results

### TDP-43 RBD binds single stranded pUG RNA

We used electrophoretic mobility shift assays (EMSAs) to investigate how TDP-43 binds pUG RNA. To prevent formation of soluble aggregates involving the CTD, we utilized constructs containing either just the RBD (amino acids 102-269), or the NTD-RBD (amino acids 1-269). Polyacrylamide gel electrophoresis (PAGE) analyses of the purified proteins and RNAs used in this study are shown in Supplementary Figure 1. To determine the binding stoichiometry and fraction of active RBD, we measured binding to (GU)_12_ RNA at high concentration of RNA (20 μM) and protein (up to 60 μM). We observed 1:1 and 2:1 protein–RNA complexes, with no further binding at higher protein concentrations (Figure 1A). These stoichiometry data indicate that ∼ 100% of TDP-43 RBD molecules are active for binding to RNA. To investigate the effect of pUG folding on RBD binding, we included K^+^ in the binding buffer and gel system and pre-folded the RNA prior to binding. The pUG fold coordinates K^+^ ions and does not fold in Li^+^ or Na^+^ buffers (40) (Supplementary Figure 2A, B). No TDP-43 binding is observed to the pUG fold, even at high concentrations (Figure 1A). These data indicate that the TDP-43 RBD can only bind to single-stranded RNA, and pUG folding prevents its ability to recognize pUG RNA. The pUG fold has very slow unfolding kinetics with a half-life of 5 days (53), resulting in an activation barrier that effectively inhibits TDP-43 binding.

**Figure 1.**
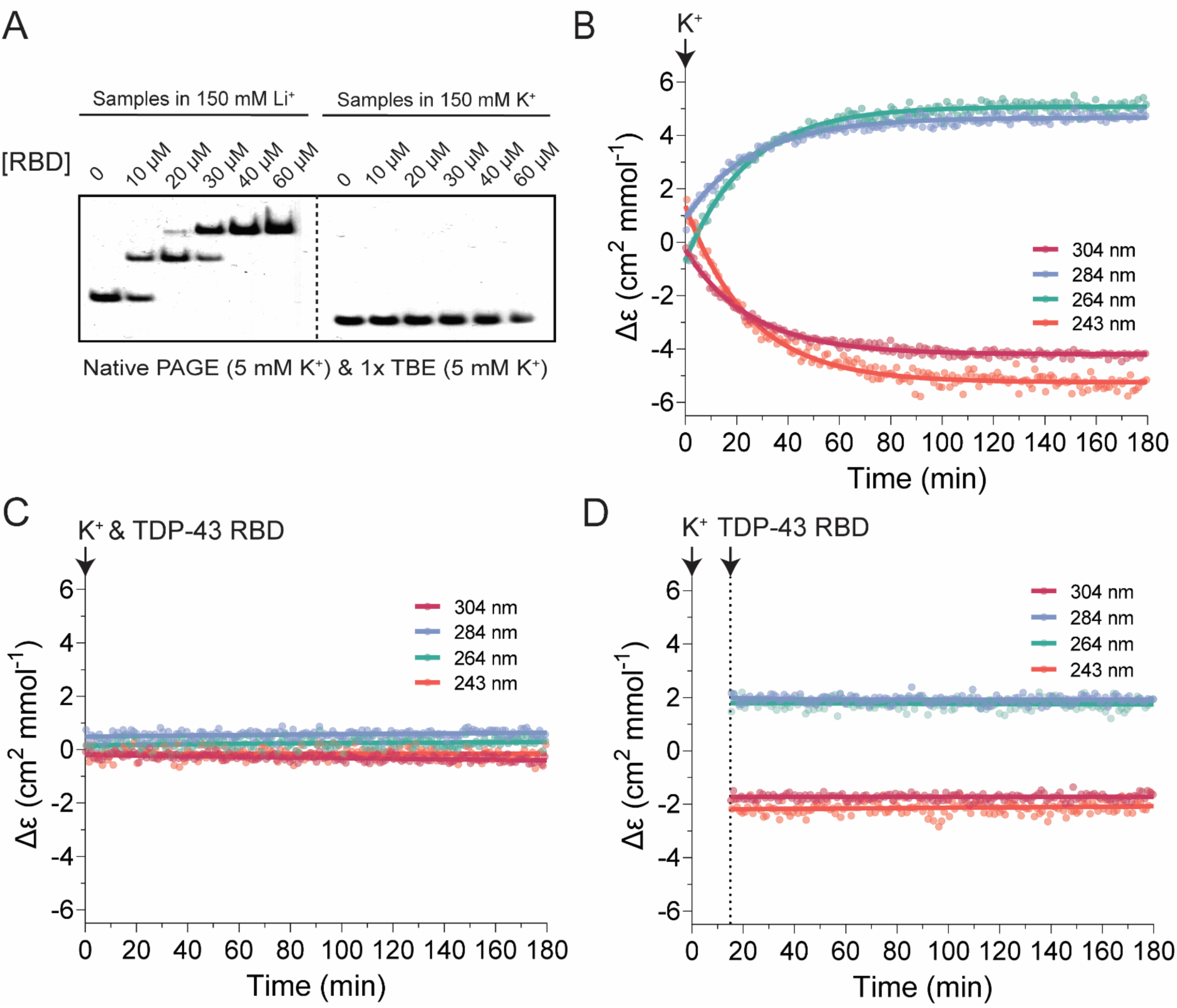
TDP-43 RBD interacts with single stranded (GU)_12_ and prevents RNA folding. (A) Native PAGE showing stoichiometric interactions between TDP-43 RBD and 20 µM (GU)_12_ RNA. RNA was pre-folded in 20 mM Bis-Tris (pH 7.0) with 150 mM Li⁺ or K⁺ before incubation with increasing concentrations of TDP-43 RBD. (B-D) CD-monitored folding of 20 µM (GU)_12_ RNA initiated by the addition of 150 mM K⁺, measured in the absence (B) or presence of 40 µM TDP-43 RBD (C, D). RBD was added at the onset of folding (C) or 15 min after initiating the reaction (D). Arrows indicate the time points at which K⁺ or TDP-43 RBD were added. CD signals at 243, 264, 284, and 304 nm, correspond to characteristic peaks of the pUG fold.

We previously used the unique circular dichroism (CD) absorption properties of the pUG fold to describe the kinetics of pUG fold formation, which is relatively slow with a half-life of ∼15 min. in 150 mM K⁺ (53). To test whether TDP-43 affects RNA folding, we used CD to monitor the kinetics of pUG RNA folding in the absence and presence of the TDP-43 RBD (Figure 1B-D). RNA folding was initiated by injection of 150 mM K⁺, and RBD was introduced either at the onset of folding or half-way through the RNA folding reaction. In the absence of protein, injection of K^+^ initiates RNA folding (pUG fold formation), as reported by the characteristic CD signals, including a negative peak at 304 nm that reports on the unique Z-form syn/anti stacking in the core of the pUG fold G4 (53) (Figure 1B). Simultaneous injection of RBD and K⁺ inhibits pUG folding, and the RNA remains unstructured for hours as indicated by a lack of RNA CD signal (Figure 1C, Supplementary Figure 3). Injection of RBD half-way through the RNA folding reaction (t=15 min.) results in 50% folded RNA after 3 h, indicating that the protein efficiently arrests the progress of the RNA folding reaction (Figure 1D, Supplementary Figure 3). These results indicate that binding of RBD efficiently prevents intramolecular pUG folding.

### The TDP-43 RBD binds pUG RNA non-cooperatively

To investigate how the RBD binds to single-stranded (GU)_12_ pUG RNA, we performed EMSA assays using 5 pM ^32^P-radiolabeled RNA, which is near the reliable detection limit for ^32^P (Figure 2A). To prevent pUG folding and ensure single stranded RNA, we used a binding buffer containing 150 mM Li^+^ instead of K^+^. Equilibrium binding conditions were demonstrated by varying incubation times up to 24 h (Figure 2A, B, Supplementary Figure 4) (54). Consistent with stoichiometry measurements at high concentrations of RNA and protein (Figure 1), two complexes were observed corresponding to one or two proteins bound to the RNA. While the first binding event forms rapidly, the second requires 24 h to reach equilibrium (Figure 2A, B, Supplementary Figure 4). At equilibrium, 5 pM protein binds nearly all the RNA in a 1:1 complex, indicating that this measurement is in the titration regime (complete binding of a limiting reagent) and the upper limit for *K_D1_* is ≤ 5 pM. Thus, the RBD binds single stranded pUG RNA with high affinity (Figure 1A).

**Figure 2.**
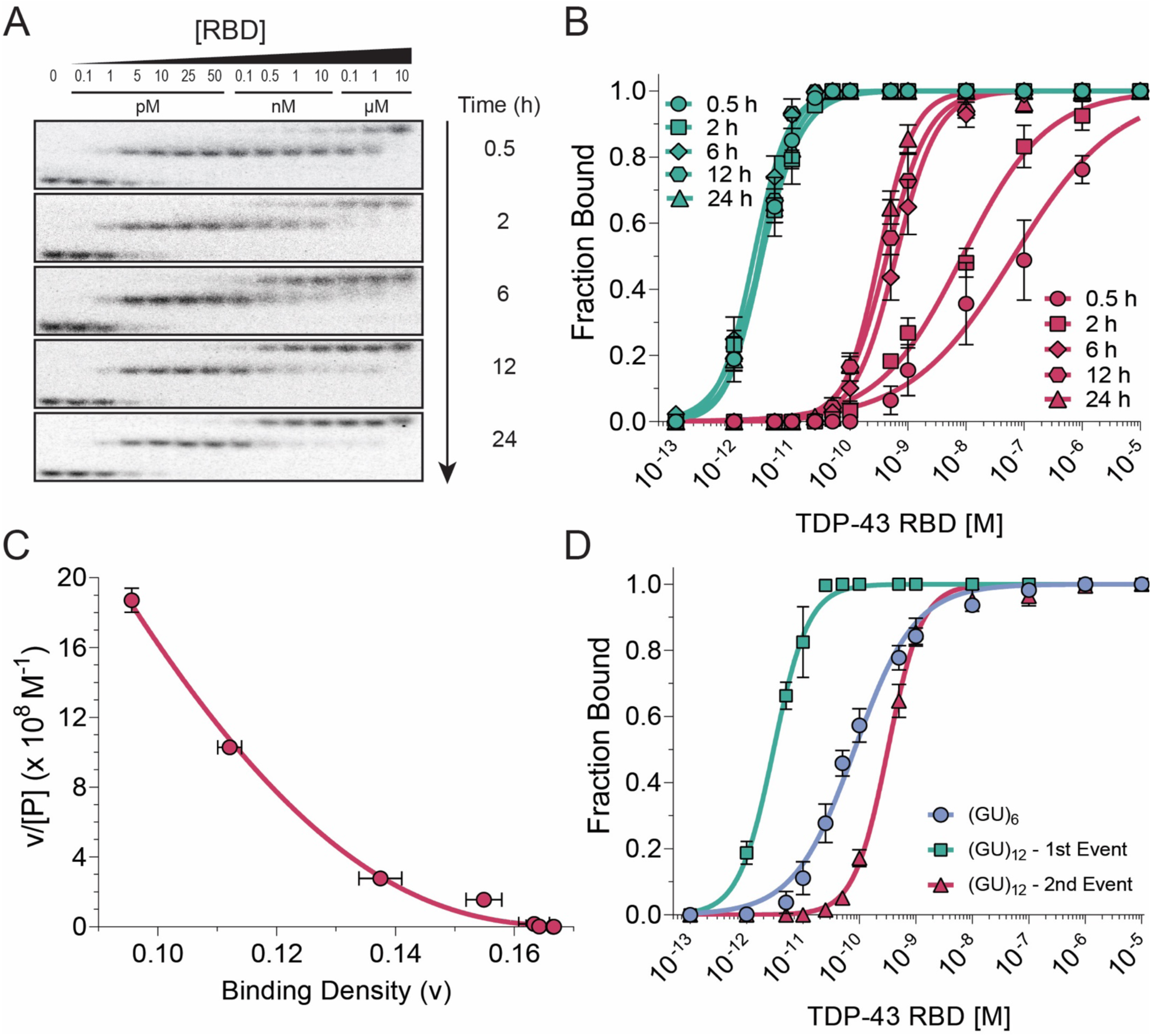
Quantitative analysis of TDP-43 RBD interaction with (GU)_12_ RNA. (A) EMSA showing TDP-43 RBD binding to 5 pM (GU)_12_ RNA after incubation for different durations (0.5-24 h) prior to electrophoresis. (B) Quantification of EMSA data. Green lines and symbols denote the first binding event; red lines and symbols denote the second. (C) Scatchard plot of the 24 h incubation data, fitted using the McGhee- von Hippel non-cooperative binding model. Binding density represents the fraction bound per dinucleotide repeat. (D) Comparison of TDP-43 RBD binding to (GU)_6_ and (GU)_12_ RNAs.

The second binding event has an apparent *K_D2_* = 318 pM (Figure 2B, Table 1). The weaker affinity for the second binding event is consistent with intrinsically anticooperative binding of the second protein and/or with intrinsically non-cooperative binding to the lattice of overlapping potential binding sites in the repetitive pUG sequence, where tight binding of the first molecule interferes with the binding of a second. To distinguish between these alternatives, we analyzed the data using a one-dimensional (1D) homogeneous lattice model (MvH model), as described by McGhee and von Hippel (55,56). The MvH model was modified to include an end correction factor to account for the finite length of the RNA, as previously described for DNA (57). This model describes the effect of overlapping binding sites and can be used to distinguish between intrinsically positive, negative and non-cooperative binding through a Scatchard-type of analysis. The MvH model also allows extraction of the intrinsic *K_D_* for binding to a single site (without the effect of overlapping sites), as well as the number of lattice units, n, covered in that binding event (Figure 2C, Table 1). The equilibrium binding data for RBD fit well to a non-cooperative MvH binding model (Figure 2C), with an intrinsic *K_D_* of 70 pM for n = 5.5 +/- 0.1 dinucleotides (di-nt) (Table 1). Therefore, the first binding event has an apparent *K_D1_* ≤ 5 pM due to the presence of multiple overlapping binding sites on the pUG RNA, which increase the fraction bound by increasing the effective concentration of binding sites. The second binding event is weaker due to overlapping sites in the 1D lattice.

**Table 1:**
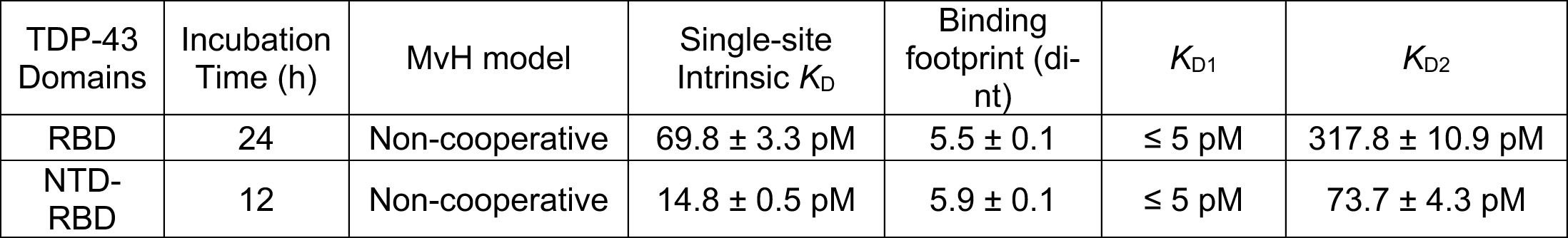
Binding constant of TDP-43 with (GU)_12_.

| TDP-43 Domains | Incubation Time (h) | MvH model | Single-site Intrinsic $K_D$ | Binding footprint (di-nt) | $K_{D1}$ | $K_{D2}$ |
| --- | --- | --- | --- | --- | --- | --- |
| RBD | 24 | Non-cooperative | $69.8 \pm 3.3$ pM | $5.5 \pm 0.1$ | $\leq 5$ pM | $317.8 \pm 10.9$ pM |
| NTD-RBD | 12 | Non-cooperative | $14.8 \pm 0.5$ pM | $5.9 \pm 0.1$ | $\leq 5$ pM | $73.7 \pm 4.3$ pM |

To further validate the binding site size, we measured binding of the RBD to the 12-nt (GU)_6_ RNA, which according to the MvH model analysis should only contain a single binding site (Supplementary Figure 5; Supplementary Table 1). Indeed, the EMSA data show single site binding with an apparent *K*_D_ of 73 pM for the RBD (Figure 2D and Supplementary Figure 5), which corresponds to the intrinsic *K*_D_ of 70 pM determined by the MvH model (Table 1). The pronounced effect of the lattice can be seen by comparing the binding of (GU)_6_ to (GU)_12_, as the first binding event to (GU)_12_ is significantly stronger than to (GU)_6_ while the second binding event is weaker (Figure 2D). We also measured the binding of RBD to (GU)_6_ in 150 mM Na^+^ buffer and obtained a nearly identical *K*_D_ of 68 pM (Figure 3). Therefore, the binding affinity is not influenced by the monovalent cation identity.

**Figure 3.**
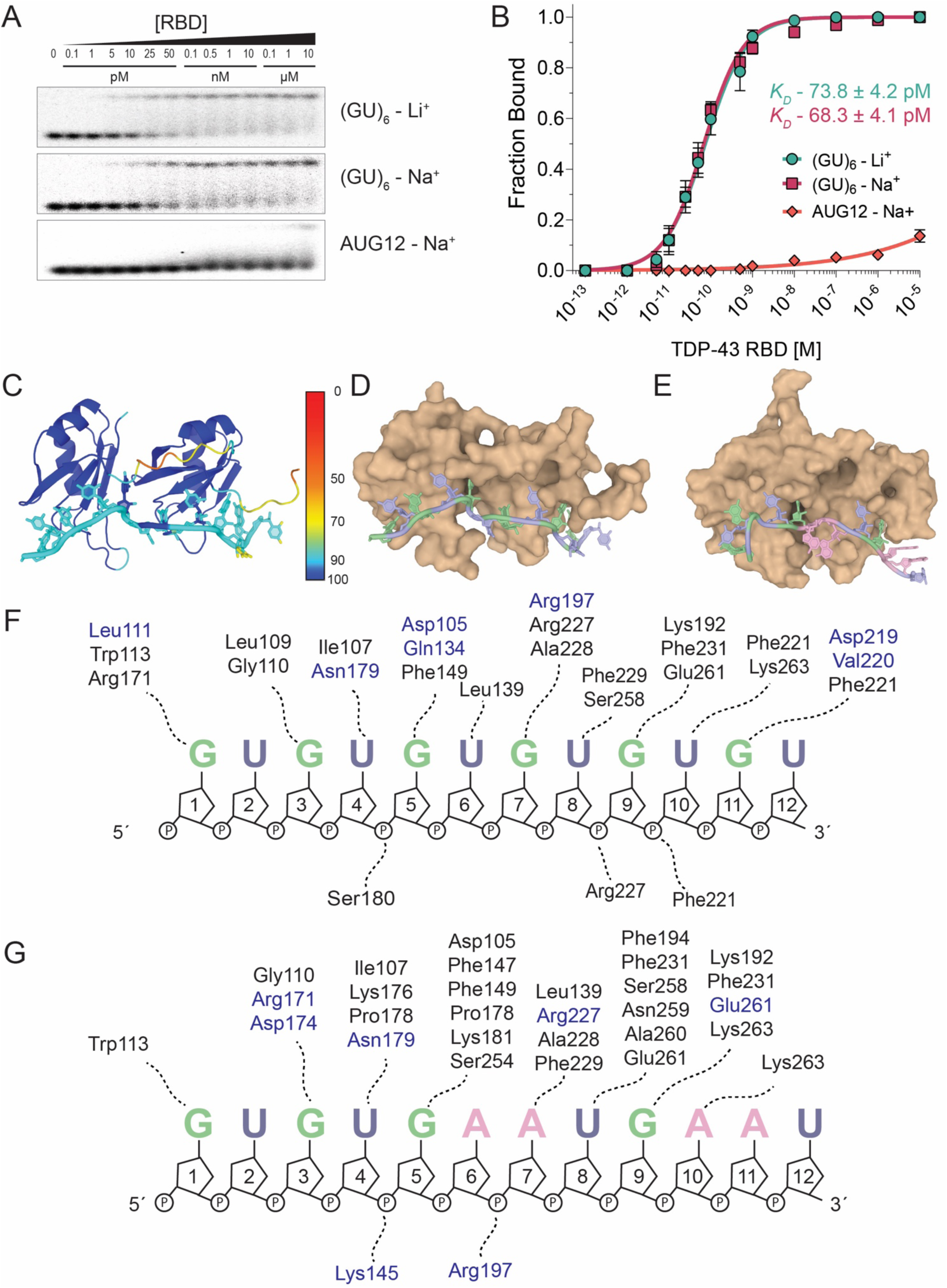
Specific Binding of TDP-43 RBD to GU repeat RNA. (A) EMSA of TDP-43 RBD binding to (GU)_6_ RNA in 150 mM Li⁺ buffer (top), (GU)_6_ RNA in 150 mM Na⁺ buffer (middle), and AUG12 RNA in 150 mM Na⁺ buffer (bottom). RNA concentrations were 5 pM for (GU)_6_ and 100 pM for AUG12 to enable detection of the complex. (B) Binding curves for (GU)_6_ in Li⁺ and Na⁺ buffers with apparent *K*_D_ values. An apparent *K*_D_ for AUG12 could not be determined. (C) Confidence scores of predicted AlphaFold3 structure of TDP-43 RBD bound to (GU)_6_ RNA. (D) Surface representations of the predicted complex. Guanosines are green and uridines are blue. (E) Surface representation of the NMR solution structure of TDP- 43 RBD bound to AUG12 RNA (PDB: 4BS2). RNA is colored as in (D) with adenosines in pink. (F) Interacting residues in the RBD-(GU)_6_ RNA complex. Hydrogen-bonding residues are shown in blue and contact residues in black. Interacting residues were identified using the PDBsum webserver. (G) Interacting residues in the RBD- AUG12 RNA complex, colored as in panel F.

### The TDP-43 RBD specifically recognizes GU repeats

To investigate the sequence specificity of the RBD, we examined binding to the 12 nt GUGUGAAUGAAU (AUG12), in which 2 UG repeats are substituted with adenosines. The AUG12 oligo was used to determine the structure of TDP-43 RBD bound to this RNA at high concentrations (0.4-0.7 mM)(35). We compared the binding affinities of the RBD for the 12mers (GU)_6_ and AUG12 (Figure 3 A, B). We were unable to measure a *K*_D_ for AUG12 and only detect weak binding at 10 μM protein (Figure 3 A, B). Thus, the RBD has a specificity for (GU)_6_ that is approximately 10^5^-fold greater than AUG12. A strong preference for binding to GU repeats is consistent with iCLIP (8) and RIP-seq data (9).

To gain further insight into the high affinity and specificity of TDP-43 for pUG RNA, we used AlphaFold 3 (58) to generate a structural model of RBD bound to (GU)_6_ (Figure 3 C, D). The resulting model is consistent with the MvH-derived footprint (Figure 3 F). Comparison with the NMR structure of RBD bound to AUG12 suggests a more continuous network of specific contacts that span nucleotides 1-11 (Figure 3 C-G). The additional sequence-specific contacts predicted by the AlphaFold model provide a potential explanation for the difference in affinity for (GU)_6_ vs. AUG12.

### The TDP-43 NTD promotes efficient multimeric assembly on long pUG RNA

We investigated the role of the NTD by analyzing the binding of NTD-RBD and comparing it to RBD. Like the RBD, two high affinity binding events to (GU)_12_ are observed at low protein concentrations (Figure 4A, B). However, we also observe a third, more slowly migrating species at the highest protein concentration tested (10 µM) (Figure 4A). This slowly migrating species is consistent with a third molecule of TDP-43 associating with two RNA-bound molecules (Supplementary Figure 6), and is also consistent with the *K*_D_ of ∼2 µM for the NTD head to tail interaction (31,32). Stoichiometric measurements performed at high concentrations indicate that the NTD-RBD is close to 100% active for binding (Supplementary Figure 6).

**Figure 4.**
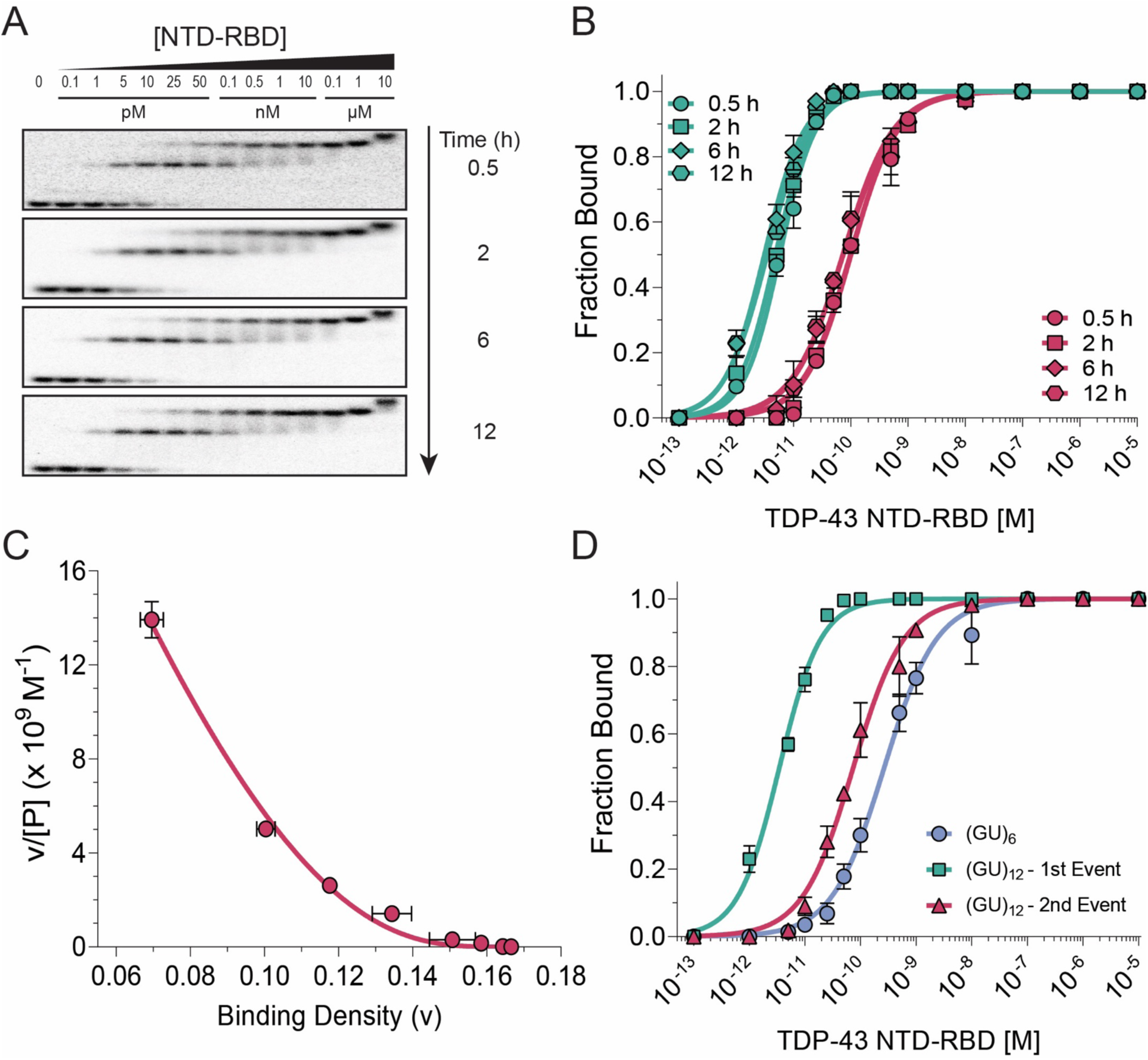
Analysis of TDP-43 NTD-RBD interaction with (GU)_12_ RNA. (A) EMSA showing TDP-43 NTD-RBD binding to 5 pM (GU)_12_ RNA after incubation for different durations (0.5- 12 h) prior to electrophoresis. (B) Quantification of EMSA data showing two distinct binding events. Green lines and symbols denote the first binding event; red lines and symbols denote the second. (C) Scatchard plot of the 12 h incubation data, fitted using the McGhee–von Hippel non-cooperative binding model. Binding density represents the fraction bound per dinucleotide repeat. (D) Comparison of TDP-43 RBD binding to (GU)_6_ and (GU)_12_ RNAs.

In contrast to RNA binding by the RBD, which takes 24 h to approach equilibrium (Figures 2A, B), the NTD-RBD approached equilibrium within 30 min for both binding events on (GU)_12_ (Figures 4A, B). As was the case for the RBD, the first binding event falls within the titration regime and has an apparent *K_D1_* ≤ 5 pM (Figure 4B, Table 1). The second binding event has an apparent *K_D2_* = 74 pM, which is significantly tighter than the second binding event for the isolated RBD (*K_D2_* = 318 pM). These results suggest that the NTD functions to efficiently recruit a second molecule onto pUG RNA, even at low (pM) concentrations, resulting in a rapid approach to equilibrium binding that cannot be achieved by the isolated RBD alone. MvH analysis of the NTD-RBD binding data (Figure 4C) reveals they fit well to a non-cooperative binding model with an intrinsic *K*_D_ of 15 +/- 1 pM, significantly stronger than the isolated RBD (intrinsic *K*_D_ = 70 +/- 3 pM; Table 1). The MvH n-value for NTD-RBD of 5.9 +/- 0.1 di-nt covered by binding one protein differs only slightly from that of the RBD (5.5 +/- 0.1 di-nt, Table 1). The NTD therefore doesn’t significantly increase the footprint of the RBD on the RNA. As with the RBD, the extremely tight first binding event of NTD-RBD on the 24-nt (GU)_12_ can be attributed to the presence of multiple overlapping binding registers. For both the RBD and NTD-RBD, the *K_D2_* values are significantly weaker than the intrinsic single site *K*_Ds_ due to non-cooperative interactions with the 1D lattice (Table 1).

We also measured binding of NTD-RBD to the single site RNA (GU)_6_, which has an apparent *K*_D_ of 251 pM (Supplementary Figure 7; Supplementary Table 1). This measurement is ∼3-fold higher than the apparent *K*_D_ of 73 pM for the isolated RBD (Supplementary Figure 5, Supplementary Table 1). Thus, the presence of the NTD has a small but measurable negative effect on binding to single sites, likely due to transient interactions between the NTD and RBD (59,60). In contrast, the intrinsic single site *K*_D_ of 15 pM for NTD-RBD binding to (GU)_12_ is 16-fold tighter than its binding to (GU)_6_ (15 vs. 251 pM). The binding curves reveal tighter binding of NTD-RBD to (GU)_12_ compared to (GU)_6_ for both binding events (Figure 4D). This contrasts with the RBD, which has a weaker second binding event (Figure 2D). The higher affinity second binding event of NTD-RBD on (GU)_12_ can be attributed to RNA facilitated protein-protein NTD interactions, which cannot occur with the RBD alone, or for the NTD-RBD on a single site (GU)_6_ RNA. The effect of protein-protein interactions is also reflected in the higher binding affinity of a second molecule of NTD-RBD vs RBD for (GU)_12_ (*K*_D2_ =74 vs 318 pM, Table 1) and a faster approach to equilibrium (0.5 h vs. 24 h). Although the NTD weakly self-associates with a *K*_D_ of ∼2 µM in the absence of RNA (31,32), RNA binding facilitates the association of a second TDP-43 molecule by creating a high local concentration of NTD, even at pM concentrations. The NTD interactions are also reflected in the MvH analysis, which shows a higher affinity intrinsic *K*_D_ for NTD-RBD vs. RBD (15 vs. 70 pM, Table 1). Despite these cooperative protein-protein interactions, NTD-RBD binding to (GU)_12_ is best described by an overall non-cooperative MvH model (Figure 4C) due to the dominant effects of overlapping high affinity binding sites within the 1D lattice. A striking property of the 1D lattice is that it leads to a highly productive initial binding event, even at concentrations below the intrinsic *K*_D_ of the protein. Dinucleotide repeats are common in human RNAs (40), but have not been previously characterized as 1D lattices.

### Facilitated dissociation of TDP-43

To further understand the TDP-43 RNA binding mechanism, we performed dissociation assays using EMSA (Figure 5A, B; Supplementary Figure 8A, B). Pre-formed RNA-protein complexes were challenged with increasing concentrations of unlabeled competitor RNA up to 10 µM, or 1000× the protein concentration of 10 nM. The 2:1 complexes resolve into free RNA with little to no 1:1 intermediates, suggesting dissociation in a concerted manner (Supplementary Figure 8A, B). This behavior is expected for the NTD-RBD, which self-associates when bound to RNA, but is also observed for the RBD. We hypothesized that this concerted dissociation may be due to end effects, as the 2:1 complexes occupy the entire length of the 24-nt (GU)_12_ RNA and place both proteins near RNA ends. To test this, we generated a 1:1 RBD:(GU)_12_ complex using a 0.5 h incubation (Figure 2A) and measured dissociation kinetics of a single RBD on the longer pUG RNA (Figure 5C, Supplementary Figure 8C). This 1:1 complex on (GU)_12_ dissociates ∼100- fold more slowly than either the 2:1 complex on (GU)_12_ or the 1:1 complex on (GU)_6_ (Figure 5A, C and Supplementary Figure 8A, C).

**Figure 5.**
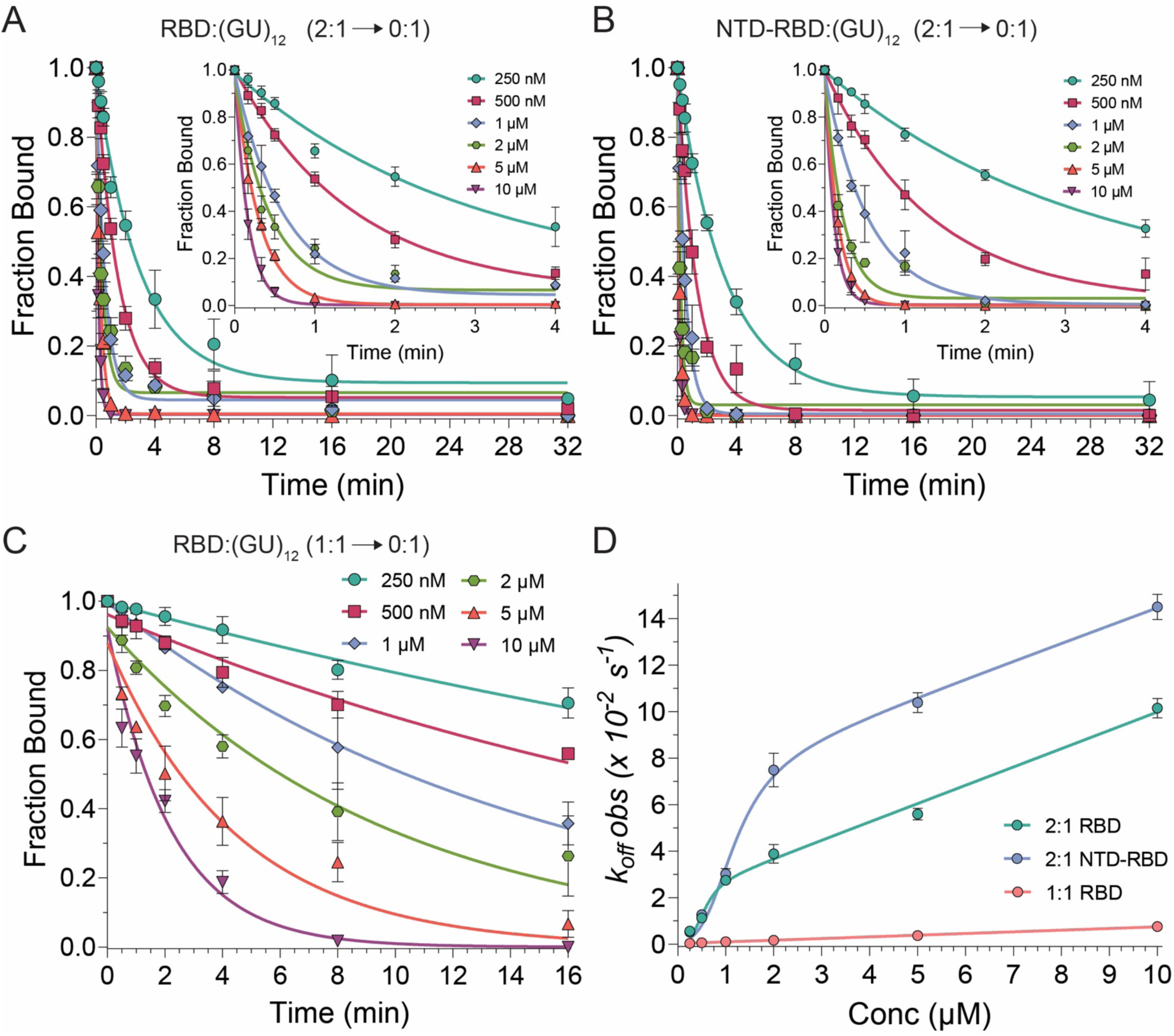
Dissociation kinetics of TDP-43 RBD and NTD-RBD complexes with (GU)_12_ RNA. (A - C) Dissociation kinetics of (A) 2:1 RBD, (B) 2:1 NTD-RBD and (C) 1:1 RBD complexes. 2:1 complexes were formed by incubating protein and RNA for 12-16 h, whereas the 1:1 complex was formed by incubation for 0.5 h. (A, B) Plots show the normalized decrease in signal intensity corresponding to the 2:1 protein-RNA bound fraction. Insets highlight the first 4 min of the dissociation regime. (C) Normalized fraction bound for the 1:1 RBD complex was fitted to a single-exponential decay to obtain the apparent rate constant (*k*_obs_). (D) Apparent dissociation rate constants (*k*_obs_) for RBD (at equilibrium and pre-equilibrium) and NTD-RBD plotted as a function of competitor RNA concentration. Data were fitted using a biphasic model that accounts for both classical and facilitated dissociation.

The observed dissociation rates exhibit biphasic behavior with an exponential phase at lower competitor RNA concentrations followed by a linear phase at higher concentrations, consistent with competitor-induced facilitated dissociation (61-68) (Figure 4D). Facilitated dissociation can occur through independent pathways that involve either occlusion of rapid rebinding events or transient formation of a ternary (or higher order complex) between TDP-43, labeled RNA, and unlabeled competitor RNA that leads to direct transfer of the protein from labeled to unlabeled RNA (Supplementary Figure 9). The data can be fit to a model that has both a saturable exponential component (classical dissociation, *k*_off_) and a linear term (exchange *k*_exch_) (63, 64). Dissociation of a 1:1 complex from (GU)_6_ RNA can also be fit to the same biphasic model (Supplementary Figure 10, Supplementary Table 2). The classical dissociation rates and exchange rates (*k_exch_*) are summarized in Table 2. Association rates were inferred from the dissociation rates and apparent *K*_D_ values and are consistent with on rates that are near the diffusion-controlled limit (Table 2). The inferred association rate for the second binding event is approximately ten-fold higher for NTD-RBD than for RBD, consistent with NTD-mediated protein-protein interactions.

**Table 2:**
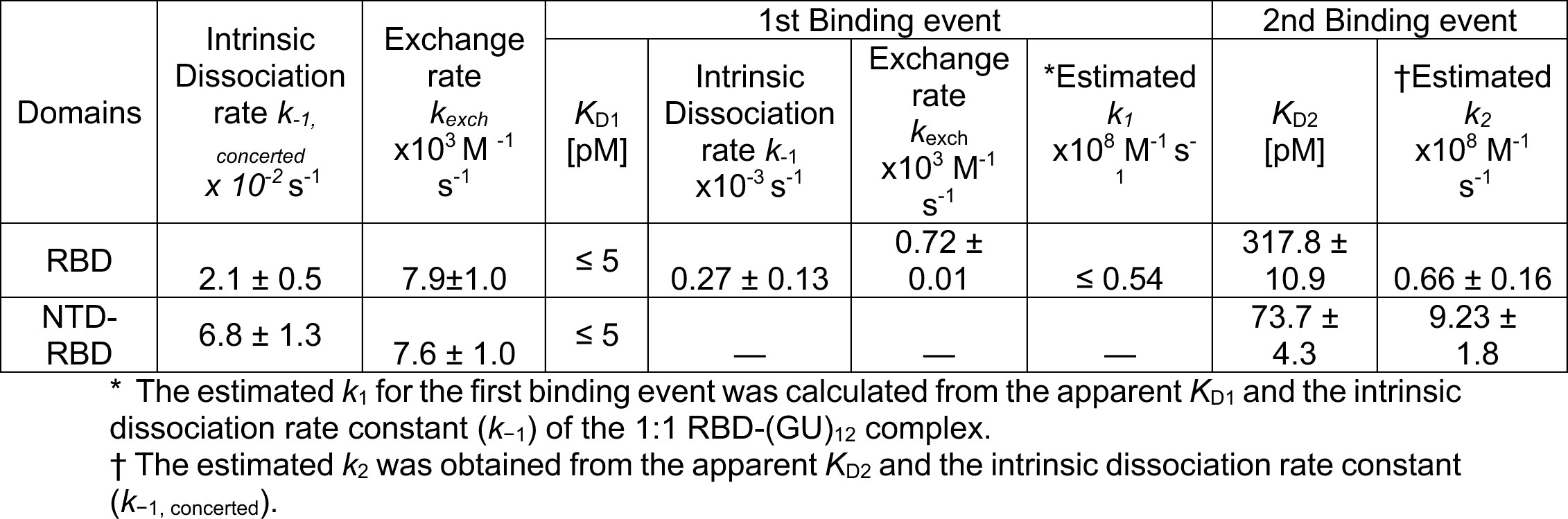
Kinetic analysis of TDP-43 RBD and NTD-RBD dissociation and exchange on (GU)_12_.

| Domains | Intrinsic Dissociation rate $k_{-1, \text{concerted}}$<br>$\times 10^{-2} \text{ s}^{-1}$ | Exchange rate $k_{\text{exch}}$<br>$\times 10^3 \text{ M}^{-1} \text{ s}^{-1}$ | 1st Binding event | | | | 2nd Binding event | |
| --- | --- | --- | --- | --- | --- | --- | --- | --- |
| | | | $K_{D1}$<br>[pM] | Intrinsic Dissociation rate $k_{-1}$<br>$\times 10^{-3} \text{ s}^{-1}$ | Exchange rate $k_{\text{exch}}$<br>$\times 10^3 \text{ M}^{-1} \text{ s}^{-1}$ | *Estimated $k_1$<br>$\times 10^8 \text{ M}^{-1} \text{ s}^{-1}$ | $K_{D2}$<br>[pM] | †Estimated $k_2$<br>$\times 10^8 \text{ M}^{-1} \text{ s}^{-1}$ |
| RBD | $2.1 \pm 0.5$ | $7.9 \pm 1.0$ | $\leq 5$ | $0.27 \pm 0.13$ | $0.72 \pm 0.01$ | $\leq 0.54$ | $317.8 \pm 10.9$ | $0.66 \pm 0.16$ |
| NTD-RBD | $6.8 \pm 1.3$ | $7.6 \pm 1.0$ | $\leq 5$ | — | — | — | $73.7 \pm 4.3$ | $9.23 \pm 1.8$ |

\* The estimated $k_1$ for the first binding event was calculated from the apparent $K_{D1}$ and the intrinsic dissociation rate constant ( $k_{-1}$ ) of the 1:1 RBD-(GU)<sub>12</sub> complex.
† The estimated $k_2$ was obtained from the apparent $K_{D2}$ and the intrinsic dissociation rate constant ( $k_{-1, \text{concerted}}$ ).

## Discussion

TDP-43 regulates RNA processing by binding to RNA and facilitating interactions with other RNA binding proteins, splicing factors, and translation machinery via an RNA-dependent mechanism (27,69). Our data indicate that TDP-43 regulates pUG RNA structure by preventing pUG folding, which is likely important for allowing additional proteins to interact with RNA. We reveal an interplay between intramolecular RNA folding and protein binding, as the soluble TDP- 43 RBD preferentially binds to single stranded pUG RNA with an affinity of at least 10^7^-fold greater than pre-folded pUG fold RNA. TDP-43 loss of function disrupts RNA processing, causing mis-splicing and the inclusion of cryptic exons in genes essential for neuronal function (8,10,14-16,18,19,70,71). We therefore hypothesize that loss of TDP-43 function may be accompanied by RNA misfolding, adversely affecting downstream interactions between RNA and other proteins.

The association rates of TDP-43 RBD and NTD-RBD are at or near the diffusion-controlled limit (Table 2). The nuclear concentration of TDP-43 has been estimated at 2.4-2.9 μM (72), and at these concentrations, TDP-43 binding to RNA is kinetically favored over pUG folding (Figures 1 and 6). This suggests an important function for TDP-43 in maintaining RNA in a single stranded state. By controlling RNA folding, TDP-43 may influence downstream RNA-protein interactions in a wide range of pathways including pre-mRNA splicing, lncRNA function, and polyadenylation. In comparison to a database analysis of protein-nucleic acid binding affinities (73), the binding data presented here show that TDP-43 ranks amongst the tightest RNA-protein interactions measured for a human protein (Supplementary Table 3).

**Figure 6.**
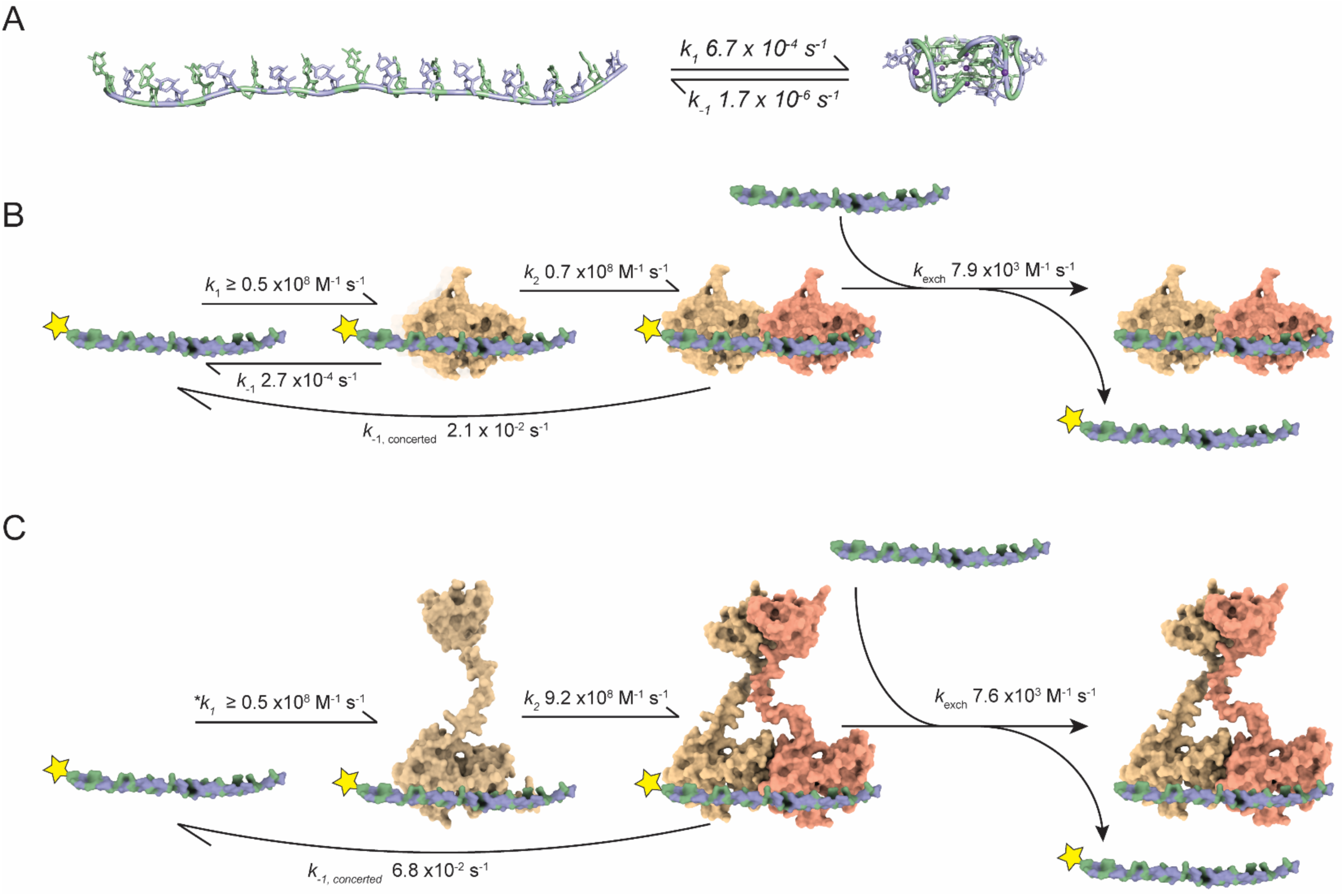
Rate constants for pUG RNA folding and TDP-43 binding. (A) Folding kinetics of (GU)_12_ RNA in the presence of 150 mM K⁺ (adapted from Petersen et al., 2025). Guanosines are green and uridines are blue, shown in ribbon and stick cartoons. The folded conformation corresponds to PDB ID 7MKT. (B) Binding kinetics of TDP-43 RBD to (GU)_12_ RNA. The yellow star denotes ^32^P labeled (GU)_12_. RNA is colored as in (A), shown in space-filling representation. (C) Binding mechanism of TDP- 43 NTD-RBD to (GU)_12_ RNA. The NTD promotes faster association of a second molecule through RNA-assisted dimerization. The asterisk (*) indicates the estimated *k*_1_ for the first association event using the dissociation rate of the 1:1 RBD-(GU)_12_ complex, because a 1:1 NTD-RBD-(GU)_12_ complex could not be isolated for measurement.

Understanding TDP-43 binding to (GU)_12_ is important as this sequence is present 20,000 times in human RNAs (40). Binding to (GU)_12_ is non-cooperative for both RBD and NTD-RBD, due to high affinity interactions with overlapping binding sites. While these overlapping sites facilitate the initial binding event, most binding events exclude a second as the remaining sites are either competitive (overlapped with occupied sites) or futile (too short to accommodate binding) (Supplementary Figure 11). The overlapping binding sites lead to long equilibration times, as the protein must sample many sites before finding the register that allows binding of a second molecule. This mechanism is well captured by the McGhee-von Hippel (MvH) 1D lattice binding model (55) and further validated by comparative binding analysis with shorter, single-site pUG RNAs. To our knowledge, this is the first time a dinucleotide repeat has been shown to behave as a 1D lattice. 1D lattices do not have to be rigid, and have been previously observed for single nucleotide repeats in flexible oligo(dT) ssDNA (74). Although flexible, single stranded sequences may form bulges or looped regions, transient formation of such structures do not appear to impact the dinucleotide lattice and are likely resolved during equilibrium binding. Hairpin secondary structures are also unlikely to impact TDP-43 recognition of pUG RNAs as nearest neighbor interactions between GU/UG wobbles are very weak (75).

Analysis of our binding data yield a binding site size of 5.5 di-nt for the RBD and 5.9 di-nt for the NTD-RBD. These data agree well with all atom models which predict a footprint of 11-12 nt (Figures 3 and 6). For any pUG RNA where *M* is the RNA length in di-nt, and *n* is the footprint of TDP-43 in di-nt, the number of overlapping sites (N) is N = *M – n + 1*. For the 12 di-nt RNA sequence (GU)_12_, there are 7 complete binding sites, and 5 out of 7 sites prevent the binding of a second molecule (Supplementary Figure 11). The apparent *K*_D1_ of ≤ 5 pM is in close agreement with the intrinsic *K*_D_, strengthened by the predicted number of binding sites (N = 7) in the dinucleotide lattice. For example, the intrinsic *K*_D_/N is 10 pM and 2 pM for the RBD and NTD-RBD, respectively (Table 1). The apparent *K*_D2_ values are also in very close agreement with the intrinsic *K*_D_ modulated by the fraction of available binding sites (N = 2/7) produced by the first binding event (Supplementary Figure 11). For the RBD, the intrinsic *K*_D_/N for the second binding event is 244 pM and the measured value of *K*_D2_ is 318 pM (Table 1). For the NTD-RBD, the intrinsic *K*_D_(N) for the second binding event is 52 pM and the measured *K*_D2_ is 74 pM (Table 1). The 1D lattice binding model also explains the very slow dissociation rate of the 1:1 complex of RBD-(GU)_12_, as the overlapping binding sites provide multiple opportunities for partial dissociation to be followed by protein reengagement in a different register (Supplementary Figure 11).

Our data show that TDP-43 can dissociate through both classical and facilitated exchange mechanisms. Classical dissociation defines the intrinsic off-rate of the protein, whereas facilitated exchange is a process in which RNA displaces bound TDP-43 either through direct transfer via ternary interactions (64,67) or by preventing rebinding after dissociation (62,76)(Figure 6 B, C, Supplementary Figure 9). The facilitated exchange of RNA on TDP-43 *in vitro* is observed as a linear exchange rate and is operational at RNA concentrations above 2 μM (Figure 4D). Facilitated exchange may be further favored in crowded cellular environments such as RNA-protein condensates.

The high affinity and specificity of TDP-43 for GU repeats reported here differ significantly from previous studies. The affinity of RBD for (GU)_6_ is >300-fold tighter than a previous report, which used classical ITC experiments to measure an apparent *K*_D_ of 23 nM in a 50 mM K^+^ buffer (35). A possible reason for this discrepancy is the fact that classical ITC experiments are not sensitive enough to measure low picomolar *K*_D_ values, due to the small amount of enthalpy generated at such low concentrations. For this reason, measurement of low pM binding affinities by ITC has required alternative, more sensitive approaches (77,78). Another study reported that TDP-43 binds to the 13-nt sequence A(GU)_6_ with a *K*_D_ of 800 pM, but did not establish equilibrium conditions or the fraction of active protein, and used an oligo concentration very close to the *K*_D_, which is in the titration regime and not an affinity measurement (21). The same issues apply to another study that reported an 8 nM *K*_D_ value for RBD binding to (UG)_6_ (45). Such errors in reported binding measurements are unfortunately common (54). We could not reproduce the previously published EMSA data for AUG12, which was previously described as having similar binding affinity as (GU)_6_ (35). We only detect very weak binding to AUG12, which differs in affinity by 5 orders of magnitude (Figure 3). Our data indicate that TDP-43 binds GU repeats with high specificity.

Our conclusion that TDP-43 binds pUG RNA non-cooperatively through lattice interactions contrasts markedly with a previous study that concluded positive cooperativity for TDP-43 RBD binding (47). In this study, DNA oligos (dGT)_6_ and (dGT)_12_ were used as proxies for RNA and binding was assessed by classical ITC performed in one direction. The DNAs were titrated into excess RBD at high concentration (14-16 μM) and 2 transitions were observed, the first fit to a *K*_D_ = 51 nM and the second to a *K*_D_ = 0.4 nM (47). Cooperativity was concluded from comparison of the two fitted *K*_D_ values. Analysis of multi-site binding interactions by ITC can be challenging and ideally require demonstration of reversibility in both the forward and reverse directions (79) or alternative analysis methods (80,81) and as previously noted, ITC is typically not sensitive enough to accurately measure pM binding events. It seems plausible that the reported *K*_D1_ and *K*_D2_ values (47) were reversed as the first measured binding event, with excess RBD in the cell and limiting DNA, likely corresponded to a 2:1, and not 1:1 RBD:DNA complex. Advantages of the EMSA method utilized here is the low picomolar sensitivity and ability to directly observe and track the stoichiometries of 1:1 and 2:1 complexes. Our data clearly show non-cooperative 2:1 binding for both RBD and NTD-RBD on (GU)_12_ RNA and are summarized in Figure 6, Tables 1 and 2, and Supplementary Tables 1 and 2.

We reveal an important role for the TDP-43 NTD in facilitating multi-site binding on pUG RNA by enhancing the binding affinity of a second molecule (by ∼4-fold) and greatly reducing the amount of time required to reach equilibrium (0.5 vs. 24 h). These observations reveal a mechanism in which TDP-43 initially binds with high affinity and can dynamically organize along pUG motifs, preventing RNA intramolecular folding. The RNA remodeling activity of TDP-43 may be important for many processes, including transcriptional regulation. DNA methyltransferase 1 (DNMT1) methylates CpG DNA sites to transcriptionally silence heterochromatin and is inhibited by pUG folds (82). By preventing accumulation of pUG folds, TDP-43 may indirectly assist in the normal function of DNMT1. DNMT1 dysfunction is linked to multiple neurodegenerative conditions (83-87). Similarly, Polycomb Repressive Complex 2 (PRC2), which silences genes involved in neurodegeneration (88), is also inhibited by RNA G4 structures, including pUGs (89-91). Thus, pUG fold accumulation could inhibit DNMT1 and PRC2 and contribute to transcriptional dysregulation in disease. Recent studies have broadly linked TDP-43 loss of function (LOF) to age-related changes in DNA methylation (92-94) and aging is a key risk factor in ALS (95,96). Together, these findings suggest that TDP-43 LOF could contribute to disease progression by leading to the accumulation of pUG folds. Since there are >20,000 pUG RNAs with 12 or more repeats in human RNAs, an open question is whether there is enough TDP-43 present to prevent the formation of all pUG folds, as this will likely depend on RNA expression levels and decay rates. For example, the large non-coding RNA NEAT1 is upregulated in many cancer cells and has 29.5 GU repeats (8,97). In the future, it will be interesting to further understand how TDP-43 remodels RNA structures in cells, and how this process is altered in diseases such as ALS, FTLD and cancer.

## Materials and Methods

RNAs used in the study:

(GU)_6_ – 5′- rGrUrGrUrGrUrGrUrGrUrGrU - 3′

(GU)_12_ – 5′- rGrUrGrUrGrUrGrUrGrUrGrUrGrUrGrUrGrUrGrUrGrUrGrU - 3′

AUG12 – 5′ - rGrUrGrUrGrArArUrGrArArU - 3′

DNAs used in the study:

(GU)_12_ template: 5′ - ACACACACACACACACACACACACTATAGTGAGTCGTATTAGAA - 3′

(GU)_12_ promoter: 5′ - TTCTAATACGACTCACTATAGTGTGTGTGTGTGTGTGTGTGTGT - 3′

AUG12 template: 5′ - ATTCATTCACACTATAGTGAGTCGTATTAGAA - 3′

AUG12 promoter: 5′ - TTCTAATACGACTCACTATAGTGTGAATGAAT - 3′

### *In vitro* transcription

Double-stranded DNA templates for *in vitro* transcription were prepared from synthetic oligonucleotides purchased from Integrated DNA Technologies, Inc. Template and promoter oligonucleotides were mixed at equimolar concentrations and annealed by heating to 95 °C for 2- 5 min, followed by cooling to room temperature for 15–20 min. AUG12 and (GU)_12_ RNAs were prepared by in vitro transcription using T7 RNA polymerase and the double-stranded DNA transcription primers described above.

### Purification of RNA

(GU)_6_ RNA was chemically synthesized by GE Healthcare Dharmacon, Inc. or Integrated DNA Technologies, Inc. All RNAs were purified on 20% denaturing polyacrylamide gels (19:1 acrylamide:bis-acrylamide, 8 M urea, 89 mM Tris, 89 mM boric acid, 2 mM EDTA). RNA bands were visualized by UV shadowing, excised, and extracted by diffusion overnight at room temperature in crush-and-soak buffer (300 mM sodium acetate, 50 mM HCl, 1 mM EDTA, pH 5.6). The extracts were filtered through a 0.2-μm Steriflip® vacuum filter (Millipore®). AUG12 and (GU)_6_ RNAs were concentrated and buffer-exchanged into nuclease-free water using 3 kDa Amicon® Ultra centrifugal filters (MilliporeSigma). Alternative purification methods, such as ethanol precipitation or Hi-Trap QFF chromatography, were avoided for these shorter RNAs because they led to significant sample loss. (GU)_12_ RNA was further purified using a 1 mL or 5 mL Hi-Trap QFF column (GE Healthcare) pre-equilibrated with 10 mM KH_2_PO_4_, 10 mM K_2_HPO_4_, 100 mM NaCl, and 1 mM EDTA. The column was washed with 5 column volumes of equilibration buffer, and RNA was eluted with the same buffer containing 2 M NaCl. Eluted RNA was concentrated using 3 kDa Amicon® Ultra filters and buffer-exchanged into nuclease-free water. Purity of the RNA used in the study is shown in Supplementary Figure 1B.

Transcribed AUG12 and (GU)_12_ were treated with Quick CIP (New England Biolabs) to remove 5′ phosphates. Reactions were incubated overnight at 37°C to ensure complete dephosphorylation, then heat-inactivated at 70°C for 15-20 minutes. The RNA was purified using Monarch^®^ RNA Cleanup Columns (New England Biolabs) per the manufacturer’s instructions, with modifications to improve recovery of short RNAs. Specifically, 300 µL of ethanol was added for (GU)_12_ and 450 µL for AUG12 prior to column binding to enhance RNA recovery.

RNA purity was confirmed by denaturing PAGE before labeling. 5′ end labeling of AUG12, (GU)_6_, and (GU)_12_ RNAs was performed using T4 Polynucleotide Kinase (New England Biolabs) and γ-³²P ATP (PerkinElmer, Inc.) following the manufacturer’s instructions with minor modifications. The PNK reaction was carried out at 37°C for 1.5 h using a PCR thermocycler, followed by heat inactivation at 70°C for 15 minutes. Labeled AUG12 and (GU)_6_ RNA was resolved on 20% denaturing PAGE, recovered via crush-and-soak, filtered, and buffer-exchanged using 3 kDa Amicon Ultra Centrifugal Filters (Millipore). Labeled (GU)_12_ was purified using Monarch^®^ RNA Cleanup Columns. The final purified RNA concentrations were determined from the total RNA concentrations in the labeling reaction, adjusted for purification yields by comparing pre- and post-purification samples by denaturing PAGE and autoradiography. All RNA samples were pre-folded by the addition of the appropriate buffer, heating the samples in 1 L of 90 °C water, and slowly cooling to room temperature for 5-6 hours.

### Protein expression and purification

A plasmid encoding human TDP-43 RBD was kindly provided by Peter Josef Lukavsky (CEITEC, Masaryk University). The construct containing the N-terminal domain fused to the RBD (NTD-RBD) was generated by inserting a codon-optimized gBlock encoding the human TDP-43 N-terminal domain (Integrated DNA Technologies) into the RBD plasmid using HiFi DNA Assembly (New England Biolabs). Both proteins were purified using the same procedure. *E. coli* BL21(DE3) pLysS cells (Invitrogen) were transformed with the expression plasmid and cultured in Luria Broth (LB) medium at 37 °C. A 2% inoculum was used to initiate secondary cultures in Terrific Broth (TB) supplemented with 1% glycerol, 100 µg/mL kanamycin, and 30 µg/mL chloramphenicol. Cultures were grown at 37 °C until reaching an OD_600_ of 1.5-2.0, chilled on ice for 30 min, and induced with 1 mM IPTG. Expression was carried out overnight at 16 °C. Cells were harvested by centrifugation and resuspended in IMAC buffer (20 mM Tris, pH 8.0, 500 mM NaCl, 20 mM imidazole, 1 mM TCEP-HCl, 10% glycerol) containing DNase I, lysozyme, 1 mM PMSF, and Protease Inhibitor Cocktail Set III (Calbiochem). Cells were lysed by sonication (15 min, 10 s on / 20 s off), and the lysate was clarified by centrifugation at 25,000 rpm for 40–60 min. The supernatant was filtered through a 0.45 µm syringe filter and loaded onto Ni-NTA agarose resin (QIAGEN). Bound protein was eluted using IMAC buffer containing 500 mM imidazole. Eluted protein was dialyzed overnight at 4 °C against ion exchange (IEX) buffer (20 mM HEPES, pH 7.0, 1 mM TCEP-HCl, 10% glycerol, 150 mM NaCl) in the presence of ∼1 mg TEV protease. A second Ni-NTA purification step (Re-Ni) may be performed but is optional. The dialyzed protein was diluted 3-5 fold with IEX buffer and loaded onto a 1 mL HiTrap Heparin HP column (Cytiva). Protein was eluted using a linear gradient from 50 to 1000 mM NaCl. Purity was assessed using Mini-PROTEAN TGX Stain-Free Gels (Bio-Rad). If a 70 kDa contaminant was detected, an optional additional purification using a 1 mL HiTrap SP HP column was performed. Fractions with an A_260_/A_280_ ≤ 0.65 were pooled and dialyzed against storage buffer (20 mM Bis-Tris, pH 7.0, 300 mM NaCl, 1 mM TCEP, 10% glycerol). Proteins were concentrated and stored at −20 °C until use. Before each experiment, proteins were dialyzed into the assay-specific buffer to ensure optimal experimental conditions.

### Circular Dichroism

CD spectra were recorded using an AVIV Model 420 CD spectrometer with a quartz cell of 1 mm path length. Wavelength scans were collected from 220 to 340 nm with a step size of 1 nm and an averaging time of 3 s per point. Measurements were performed at 25 °C, and buffer spectra were subtracted. The data were converted to molecular CD absorption (Δε) using the equation:

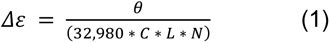

In this equation, *θ* is the observed CD signal (in millidegrees), *C* is the RNA concentration (in molar units, M), *L* is the cuvette path length (in centimeters), and *N* is the number of nucleotides in the RNA molecule.

For kinetic experiments, folding was monitored at four wavelengths characteristic of the pUG fold (243, 264, 284, and 304 nm), with 340 nm as a reference. Folding was initiated by manual mixing, and the dead time was accounted for before data collection. Ellipticity was recorded for 3 h with 180 data points, using a 3 s averaging time per point. The raw CD signal was converted to molar ellipticity (Δε) using the equation shown above. For experiments involving protein, the protein-only signal was measured and subtracted to obtain the RNA spectrum. Each kinetic experiment was performed in duplicate to ensure reproducibility. Folding half-lives were obtained by fitting Δε vs time data to a single-exponential model using least squares regression in GraphPad Prism v10. Protein contributions to the CD signal were subtracted before analysis.

### EMSA analysis

Binding reactions were performed by incubating 5 pM ^32^P-labeled RNA with increasing concentrations of protein. Reactions were carried out in buffer containing 20 mM BIS-TRIS (pH 7.0), 150 mM LiCl, 1 mM TCEP-HCl, 1 mM EDTA, 20% sucrose (w/v), 0.01% NP-40 (v/v), 0.01 mg/mL sodium heparin, and 0.01 mg/mL BSA. Samples (20 µL) were incubated at room temperature for the times specified in the corresponding figures before loading onto gels. Gels were 8% polyacrylamide (29:1 acrylamide:bis-acrylamide, 89 mM Tris-borate, 2 mM EDTA, pH 8.0). All gels were pre-run at 5W for at least 30 mins before sample loading. Electrophoresis was run at 5 W for 2 h for (GU)_6_ and AUG12, and 2.5 h for (GU)_12_ at 4 °C.

Gels were exposed to phosphorimaging screens for at least 21 h and imaged using an Amersham™ Typhoon™ 5 biomolecular imager (Cytiva) at 100 µm resolution with the highest sensitivity (S4000). Gel images were processed in Fiji. Contrast was enhanced by normalizing saturated pixels, and smoothing was applied to reduce noise. Two background subtraction steps were applied to ensure accurate quantification. The first step enhanced band definition relative to the surrounding gel. The second step isolated the signal from the bands by subtracting the background measured in adjacent regions. Only clearly resolved bands corresponding to bound and free RNA were quantified. For longer incubation times, decreased bound complex intensity was carefully verified across all lanes to ensure accurate fraction bound. All samples were processed under identical conditions. Binding data were quantified using GraphPad Prism v10. Apparent dissociation constants (*K*_D_) were estimated by fitting the binding curves to a standard sigmoidal model available in Prism. A previously described (57) form of the McGhee-von Hippel equation with an end-correction factor was used in this study:

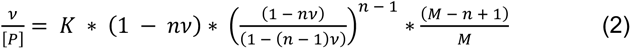

Here, *v* represents the binding density (calculated as *fraction bound divided by the number of di- nucleotide repeats in the RNA*), [P] is the free protein concentration in solution, *K* is the intrinsic association constant, n is the binding footprint of the protein (number of dinucleotides occupied per molecule), and M is the total number of dinucleotides in the RNA.

### AlphaFold3 structure prediction

AlphaFold3 was employed to predict structures of the TDP-43 NTD-RBD dimer and of the TDP- 43 NTD-RBD in complex with (GU)_6_. Models were rendered in PyMOL for further analysis and figure preparation.

### Dissociation measurements

Dissociation experiments were performed using pre-formed complexes containing 10 nM TDP-43 protein and 10 pM radiolabeled RNA. Complexes were challenged with increasing concentrations of unlabeled competitor RNA (500 nM–10 µM final). All reactions were carried out in 20 mM BIS-TRIS (pH 7.0), 150 mM LiCl, 1 mM TCEP, 1 mM EDTA, 0.01% NP-40 (v/v), 20% sucrose (w/v), 0.01 mg/mL BSA, and 0.01 mg/mL heparin. Equal volumes were used for all experiments. For zero-time controls, 10 µL of pre-formed complex was mixed with 10 µL of buffer. Dissociation reactions were initiated by mixing 10 µL of pre-formed complex with 10 µL of competitor RNA solution, giving final concentrations of 10 nM protein, 10 pM radiolabeled RNA, and 500 nM–10 µM competitor RNA. Preformed (GU)_6_ complexes were incubated for 2-4 h, as longer incubation reduced complex intensity. (GU)_12_ complexes were incubated for 12 h to reach near-equilibrium. Pre-equilibrium (GU)_12_ complexes were prepared by incubating for 30 min. Aliquots were removed at defined time points and immediately loaded onto a pre-running 8% native polyacrylamide gel (1× TBE, 4 °C). Gels were run under non-denaturing conditions and analyzed as described for EMSA experiments. Dissociation kinetics were analyzed by fitting data to a single-exponential decay model in GraphPad Prism v10 to determine apparent rate constants (*k*_obs_). A previously described biphasic model (63, 64) was used to account for both classical dissociation and competitor-facilitated RNA exchange.

## Acknowledgements

A plasmid encoding human TDP-43 RBD was kindly provided by Peter Josef Lukavsky (CEITEC, Masaryk University). Circular Dichroism data were obtained at the University of Wisconsin-Madison Biophysics Instrumentation Facility, which was established with support from the University of Wisconsin-Madison and grants BIR-9512577 (NSF) and S10RR013790 (NIH). This study was supported by NIH grants R35 GM118131 to S.E.B. and R35 GM136261 to A.A.H.

## Author Contributions

R.V.: Conceptualization, data curation, formal analysis, investigation, methodology, visualization, validation, writing- original draft preparation and review and editing. T.K.: Formal Analysis, Investigation, Methodology, Validation, Writing – Review & Editing, S.R.: Investigation, Methodology, M.T.R.: formal analysis, methodology, writing- review and editing, A.A.H.: funding acquisition, resources, supervision, writing- review and editing, S.E.B.: Conceptualization, data curation, formal analysis, funding acquisition, methodology, project administration, resources, supervision, visualization, validation, writing- original draft preparation and review and editing.

## References

1. Neumann, M., Sampathu, D.M., Kwong, L.K., Truax, A.C., Micsenyi, M.C., Chou, T.T., Bruce, J., Schuck, T., Grossman, M., Clark, C.M. et al. (2006) Ubiquitinated TDP-43 in frontotemporal lobar degeneration and amyotrophic lateral sclerosis. Science, 314, 130–133.

2. Sreedharan, J., Blair, I.P., Tripathi, V.B., Hu, X., Vance, C., Rogelj, B., Ackerley, S., Durnall, J.C., Williams, K.L., Buratti, E. et al. (2008) TDP-43 mutations in familial and sporadic amyotrophic lateral sclerosis. Science, 319, 1668–1672.

3. Balendra, R., Sreedharan, J., Hallegger, M., Luisier, R., Lashuel, H.A., Gregory, J.M. and Patani, R. (2025) Amyotrophic lateral sclerosis caused by TARDBP mutations: from genetics to TDP-43 proteinopathy. Lancet Neurol, 24, 456–470.

4. Cairns, N.J., Neumann, M., Bigio, E.H., Holm, I.E., Troost, D., Hatanpaa, K.J., Foong, C., White, C.L., 3rd, Schneider, J.A., Kretzschmar, H.A. et al. (2007) TDP-43 in familial and sporadic frontotemporal lobar degeneration with ubiquitin inclusions. Am J Pathol, 171, 227–240.

5. Josephs, K.A., Whitwell, J.L., Weigand, S.D., Murray, M.E., Tosakulwong, N., Liesinger, A.M., Petrucelli, L., Senjem, M.L., Knopman, D.S., Boeve, B.F. et al. (2014) TDP-43 is a key player in the clinical features associated with Alzheimer’s disease. Acta Neuropathol, 127, 811–824.

6. Josephs, K.A., Murray, M.E., Whitwell, J.L., Tosakulwong, N., Weigand, S.D., Petrucelli, L., Liesinger, A.M., Petersen, R.C., Parisi, J.E. and Dickson, D.W. (2016) Updated TDP- 43 in Alzheimer’s disease staging scheme. Acta Neuropathol, 131, 571–585.

7. Xiao, S., Sanelli, T., Dib, S., Sheps, D., Findlater, J., Bilbao, J., Keith, J., Zinman, L., Rogaeva, E. and Robertson, J. (2011) RNA targets of TDP-43 identified by UV-CLIP are deregulated in ALS. Mol Cell Neurosci, 47, 167–180.

8. Tollervey, J.R., Curk, T., Rogelj, B., Briese, M., Cereda, M., Kayikci, M., König, J., Hortobágyi, T., Nishimura, A.L., Zupunski, V. et al. (2011) Characterizing the RNA targets and position-dependent splicing regulation by TDP-43. Nat Neurosci, 14, 452–458.

9. Sephton, C.F., Cenik, C., Kucukural, A., Dammer, E.B., Cenik, B., Han, Y., Dewey, C.M., Roth, F.P., Herz, J., Peng, J. et al. (2011) Identification of neuronal RNA targets of TDP- 43-containing ribonucleoprotein complexes. J Biol Chem, 286, 1204–1215.

10. Polymenidou, M., Lagier-Tourenne, C., Hutt, K.R., Huelga, S.C., Moran, J., Liang, T.Y., Ling, S.C., Sun, E., Wancewicz, E., Mazur, C. et al. (2011) Long pre-mRNA depletion and RNA missplicing contribute to neuronal vulnerability from loss of TDP-43. Nat Neurosci, 14, 459–468.

11. Buratti, E., Dörk, T., Zuccato, E., Pagani, F., Romano, M. and Baralle, F.E. (2001) Nuclear factor TDP-43 and SR proteins promote in vitro and in vivo CFTR exon 9 skipping. Embo j, 20, 1774–1784.

12. Passoni, M., De Conti, L., Baralle, M. and Buratti, E. (2012) UG repeats/TDP-43 interactions near 5’ splice sites exert unpredictable effects on splicing modulation. J Mol Biol, 415, 46–60.

13. Fiesel, F.C., Weber, S.S., Supper, J., Zell, A. and Kahle, P.J. (2012) TDP-43 regulates global translational yield by splicing of exon junction complex component SKAR. Nucleic Acids Res, 40, 2668–2682.

14. Ling, J.P., Pletnikova, O., Troncoso, J.C. and Wong, P.C. (2015) TDP-43 repression of nonconserved cryptic exons is compromised in ALS-FTD. Science, 349, 650–655.

15. Jeong, Y.H., Ling, J.P., Lin, S.Z., Donde, A.N., Braunstein, K.E., Majounie, E., Traynor, B.J., LaClair, K.D., Lloyd, T.E. and Wong, P.C. (2017) Tdp-43 cryptic exons are highly variable between cell types. Mol Neurodegener, 12, 13.

16. Humphrey, J., Emmett, W., Fratta, P., Isaacs, A.M. and Plagnol, V. (2017) Quantitative analysis of cryptic splicing associated with TDP-43 depletion. BMC Med Genomics, 10, 38.

17. Fratta, P., Sivakumar, P., Humphrey, J., Lo, K., Ricketts, T., Oliveira, H., Brito-Armas, J.M., Kalmar, B., Ule, A., Yu, Y. et al. (2018) Mice with endogenous TDP-43 mutations exhibit gain of splicing function and characteristics of amyotrophic lateral sclerosis. Embo j, 37.

18. Ma, X.R., Prudencio, M., Koike, Y., Vatsavayai, S.C., Kim, G., Harbinski, F., Briner, A., Rodriguez, C.M., Guo, C., Akiyama, T. et al. (2022) TDP-43 represses cryptic exon inclusion in the FTD-ALS gene UNC13A. Nature, 603, 124–130.

19. Brown, A.L., Wilkins, O.G., Keuss, M.J., Kargbo-Hill, S.E., Zanovello, M., Lee, W.C., Bampton, A., Lee, F.C.Y., Masino, L., Qi, Y.A. et al. (2022) TDP-43 loss and ALS-risk SNPs drive mis-splicing and depletion of UNC13A. Nature, 603, 131–137.

20. Hallegger, M., Chakrabarti, A.M., Lee, F.C.Y., Lee, B.L., Amalietti, A.G., Odeh, H.M., Copley, K.E., Rubien, J.D., Portz, B., Kuret, K. et al. (2021) TDP-43 condensation properties specify its RNA-binding and regulatory repertoire. Cell, 184, 4680–4696.e4622.

21. Grese, Z.R., Bastos, A.C., Mamede, L.D., French, R.L., Miller, T.M. and Ayala, Y.M. (2021) Specific RNA interactions promote TDP-43 multivalent phase separation and maintain liquid properties. EMBO Rep, 22, e53632.

22. Zeng, Y., Lovchykova, A., Akiyama, T., Rayner, S.L., Maheswari Jawahar, V., Liu, C., Sianto, O., Guo, C., Calliari, A., Prudencio, M. et al. (2025) TDP-43 nuclear loss in FTD/ALS causes widespread alternative polyadenylation changes. Nat Neurosci.

23. Bryce-Smith, S., Brown, A.L., Chien, M., Dattilo, D., Mehta, P.R., Mattedi, F., Barattucci, S., Mikheenko, A., Zanovello, M., Pellegrini, F. et al. (2025) TDP-43 loss induces cryptic polyadenylation in ALS/FTD. Nat Neurosci.

24. Duan, L., Zaepfel, B.L., Aksenova, V., Dasso, M., Rothstein, J.D., Kalab, P. and Hayes, L.R. (2022) Nuclear RNA binding regulates TDP-43 nuclear localization and passive nuclear export. Cell Rep, 40, 111106.

25. Zhang, X., Das, T., Chao, T.F., Trinh, V., Carmen-Orozco, R.P., Ling, J.P., Kalab, P. and Hayes, L.R. (2024) Multivalent GU-rich oligonucleotides sequester TDP-43 in the nucleus by inducing high molecular weight RNP complexes. iScience, 27, 110109.

26. Dos Passos, P.M., Hemamali, E.H., Mamede, L.D., Hayes, L.R. and Ayala, Y.M. (2024) RNA-mediated ribonucleoprotein assembly controls TDP-43 nuclear retention. PLoS Biol, 22, e3002527.

27. Freibaum, B.D., Chitta, R.K., High, A.A. and Taylor, J.P. (2010) Global analysis of TDP- 43 interacting proteins reveals strong association with RNA splicing and translation machinery. J Proteome Res, 9, 1104–1120.

28. Cheng, F., Chapman, T., Venturato, J., Davidson, J.M., Polido, S.A., Rosa-Fernandes, L., San Gil, R., Suddull, H.J., Zhang, S., Macaslam, C.Y. et al. (2025) Proteomics Analysis of the TDP-43 Interactome in Cellular Models of ALS Pathogenesis. J Neurochem, 169, e70079.

29. Shiina, Y., Arima, K., Tabunoki, H. and Satoh, J. (2010) TDP-43 dimerizes in human cells in culture. Cell Mol Neurobiol, 30, 641–652.

30. Afroz, T., Hock, E.M., Ernst, P., Foglieni, C., Jambeau, M., Gilhespy, L.A.B., Laferriere, F., Maniecka, Z., Plückthun, A., Mittl, P. et al. (2017) Functional and dynamic polymerization of the ALS-linked protein TDP-43 antagonizes its pathologic aggregation. Nat Commun, 8, 45.

31. Tsoi, P.S., Choi, K.J., Leonard, P.G., Sizovs, A., Moosa, M.M., MacKenzie, K.R., Ferreon, J.C. and Ferreon, A.C.M. (2017) The N-Terminal Domain of ALS-Linked TDP- 43 Assembles without Misfolding. Angew Chem Int Ed Engl, 56, 12590–12593.

32. Herrera, M.G., Amundarain, M.J. and Santos, J. (2023) Biophysical evaluation of the oligomerization and conformational properties of the N-terminal domain of TDP-43. Arch Biochem Biophys, 737, 109533.

33. Wright, G.S.A., Watanabe, T.F., Amporndanai, K., Plotkin, S.S., Cashman, N.R., Antonyuk, S.V. and Hasnain, S.S. (2020) Purification and Structural Characterization of Aggregation-Prone Human TDP-43 Involved in Neurodegenerative Diseases. iScience, 23, 101159.

34. Wang, A., Conicella, A.E., Schmidt, H.B., Martin, E.W., Rhoads, S.N., Reeb, A.N., Nourse, A., Ramirez Montero, D., Ryan, V.H., Rohatgi, R. et al. (2018) A single N-terminal phosphomimic disrupts TDP-43 polymerization, phase separation, and RNA splicing. Embo j, 37.

35. Lukavsky, P.J., Daujotyte, D., Tollervey, J.R., Ule, J., Stuani, C., Buratti, E., Baralle, F.E., Damberger, F.F. and Allain, F.H. (2013) Molecular basis of UG-rich RNA recognition by the human splicing factor TDP-43. Nat Struct Mol Biol, 20, 1443–1449.

36. Li, Q., Babinchak, W.M. and Surewicz, W.K. (2021) Cryo-EM structure of amyloid fibrils formed by the entire low complexity domain of TDP-43. Nat Commun, 12, 1620.

37. Sharma, K., Stockert, F., Shenoy, J., Berbon, M., Abdul-Shukkoor, M.B., Habenstein, B., Loquet, A., Schmidt, M. and Fandrich, M. (2024) Cryo-EM observation of the amyloid key structure of polymorphic TDP-43 amyloid fibrils. Nat Commun, 15, 486.

38. Cao, Q., Boyer, D.R., Sawaya, M.R., Ge, P. and Eisenberg, D.S. (2019) Cryo-EM structures of four polymorphic TDP-43 amyloid cores. Nat Struct Mol Biol, 26, 619–627.

39. Arseni, D., Hasegawa, M., Murzin, A.G., Kametani, F., Arai, M., Yoshida, M. and Ryskeldi-Falcon, B. (2022) Structure of pathological TDP-43 filaments from ALS with FTLD. Nature, 601, 139–143.

40. Roschdi, S., Yan, J., Nomura, Y., Escobar, C.A., Petersen, R.J., Bingman, C.A., Tonelli, M., Vivek, R., Montemayor, E.J., Wickens, M. et al. (2022) An atypical RNA quadruplex marks RNAs as vectors for gene silencing. Nat Struct Mol Biol, 29, 1113–1121.

41. Butcher, S.E. (2024) A left-handed RNA quadruplex directs gene silencing. Trends Biochem Sci, 49, 387–390.

42. Escobar, C.A., Petersen, R.J., Tonelli, M., Fan, L., Henzler-Wildman, K.A. and Butcher, S.E. (2023) Solution Structure of Poly(UG) RNA. J Mol Biol, 435, 168340.

43. Shukla, A., Yan, J., Pagano, D.J., Dodson, A.E., Fei, Y., Gorham, J., Seidman, J.G., Wickens, M. and Kennedy, S. (2020) poly(UG)-tailed RNAs in genome protection and epigenetic inheritance. Nature, 582, 283–288.

44. Kuo, P.H., Doudeva, L.G., Wang, Y.T., Shen, C.K. and Yuan, H.S. (2009) Structural insights into TDP-43 in nucleic-acid binding and domain interactions. Nucleic Acids Res, 37, 1799–1808.

45. Bhardwaj, A., Myers, M.P., Buratti, E. and Baralle, F.E. (2013) Characterizing TDP-43 interaction with its RNA targets. Nucleic Acids Res, 41, 5062–5074.

46. Kuo, P.H., Chiang, C.H., Wang, Y.T., Doudeva, L.G. and Yuan, H.S. (2014) The crystal structure of TDP-43 RRM1-DNA complex reveals the specific recognition for UG- and TG-rich nucleic acids. Nucleic Acids Res, 42, 4712–4722.

47. Rengifo-Gonzalez, J.C., El Hage, K., Clément, M.J., Steiner, E., Joshi, V., Craveur, P., Durand, D., Pastré, D. and Bouhss, A. (2021) The cooperative binding of TDP-43 to GU-rich RNA repeats antagonizes TDP-43 aggregation. Elife, 10.

48. Flores, B.N., Li, X., Malik, A.M., Martinez, J., Beg, A.A. and Barmada, S.J. (2019) An Intramolecular Salt Bridge Linking TDP43 RNA Binding, Protein Stability, and TDP43- Dependent Neurodegeneration. Cell Rep, 27, 1133–1150.e1138.

49. Ayala, Y.M., Pantano, S., D’Ambrogio, A., Buratti, E., Brindisi, A., Marchetti, C., Romano, M. and Baralle, F.E. (2005) Human, Drosophila, and C.elegans TDP43: nucleic acid binding properties and splicing regulatory function. J Mol Biol, 348, 575–588.

50. Ishiguro, A. and Ishihama, A. (2023) ALS-linked TDP-43 mutations interfere with the recruitment of RNA recognition motifs to G-quadruplex RNA. Sci Rep, 13, 5982.

51. Zhao, J., Yang, F., Zhang, Y., Wang, H. and Kwok, C.K. (2025) TDP-43 binds to RNA G-quadruplex structure and regulates mRNA stability and translation. Nucleic Acids Res, 53.

52. Oldani, E.G., Reynolds Caicedo, K.M., Spaeth Herda, M.E., Sachs, A.H., Chapman, E.G., Kumar, S., Linseman, D.A. and Horowitz, S. (2025) The effect of G-quadruplexes on TDP43 condensation, distribution, and toxicity. Structure, 33, 1294–1303 e1295.

53. Petersen, R.J., Vivek, R., Tonelli, M., Roschdi, S. and Butcher, S.E. (2025) The structure, folding kinetics, and dynamics of long poly(UG) RNA. Nucleic Acids Res, 53.

54. Jarmoskaite, I., AlSadhan, I., Vaidyanathan, P.P. and Herschlag, D. (2020) How to measure and evaluate binding affinities. Elife, 9.

55. McGhee, J.D. and von Hippel, P.H. (1974) Theoretical aspects of DNA-protein interactions: co-operative and non-co-operative binding of large ligands to a one-dimensional homogeneous lattice. J Mol Biol, 86, 469–489.

56. Kowalczykowski, S.C., Paul, L.S., Lonberg, N., Newport, J.W., McSwiggen, J.A. and von Hippel, P.H. (1986) Cooperative and non-cooperative binding of protein ligands to nucleic acid lattices: experimental approaches to the determination of thermodynamic parameters. Biochemistry, 25, 1226–1240.

57. Tsodikov, O.V., Holbrook, J.A., Shkel, I.A. and Record, M.T., Jr. (2001) Analytic binding isotherms describing competitive interactions of a protein ligand with specific and nonspecific sites on the same DNA oligomer. Biophys J, 81, 1960–1969.

58. Abramson, J., Adler, J., Dunger, J., Evans, R., Green, T., Pritzel, A., Ronneberger, O., Willmore, L., Ballard, A.J., Bambrick, J. et al. (2024) Accurate structure prediction of biomolecular interactions with AlphaFold 3. Nature, 630, 493–500.

59. Scott, D.D., Jena, L., Rajaram, A., Ang, J., Perez-Miller, S., Kumirov, V., Khanna, R. and Khanna, M. (2025) Identifying interactions between TDP-43’s N-terminal and RNA-binding domains. Protein Sci, 34, e70295.

60. Mohanty, P., Rizuan, A., Kim, Y.C., Fawzi, N.L. and Mittal, J. (2024) A complex network of interdomain interactions underlies the conformational ensemble of monomeric TDP-43 and modulates its phase behavior. Protein Sci, 33, e4891.

61. Erbas, A. and Marko, J.F. (2019) How do DNA-bound proteins leave their binding sites? The role of facilitated dissociation. Curr Opin Chem Biol, 53, 118–124.

62. Graham, J.S., Johnson, R.C. and Marko, J.F. (2011) Concentration-dependent exchange accelerates turnover of proteins bound to double-stranded DNA. Nucleic Acids Res, 39, 2249–2259.

63. Hadizadeh, N., Johnson, R.C. and Marko, J.F. (2016) Facilitated Dissociation of a Nucleoid Protein from the Bacterial Chromosome. J Bacteriol, 198, 1735–1742.

64. Hemphill, W.O., Fenske, R., Gooding, A.R. and Cech, T.R. (2023) PRC2 direct transfer from G-quadruplex RNA to dsDNA has implications for RNA-binding chromatin modifiers. Proc Natl Acad Sci U S A, 120, e2220528120.

65. Hemphill, W.O., Voong, C.K., Fenske, R., Goodrich, J.A. and Cech, T.R. (2023) Multiple RNA- and DNA-binding proteins exhibit direct transfer of polynucleotides with implications for target-site search. Proc Natl Acad Sci U S A, 120, e2220537120.

66. Kamar, R.I., Banigan, E.J., Erbas, A., Giuntoli, R.D., Olvera de la Cruz, M., Johnson, R.C. and Marko, J.F. (2017) Facilitated dissociation of transcription factors from single DNA binding sites. Proc Natl Acad Sci U S A, 114, E3251–E3257.

67. Kozlov, A.G. and Lohman, T.M. (2002) Kinetic mechanism of direct transfer of Escherichia coli SSB tetramers between single-stranded DNA molecules. Biochemistry, 41, 11611–11627.

68. Paul, T., Lee, I.R., Pangeni, S., Rashid, F., Yang, O., Antony, E., Berger, J.M., Myong, S. and Ha, T. (2025) Mechanistic insights into direct DNA and RNA strand transfer and dynamic protein exchange of SSB and RPA. Nucleic Acids Res, 53.

69. D’Ambrogio, A., Buratti, E., Stuani, C., Guarnaccia, C., Romano, M., Ayala, Y.M. and Baralle, F.E. (2009) Functional mapping of the interaction between TDP-43 and hnRNP A2 in vivo. Nucleic Acids Res, 37, 4116–4126.

70. Estades Ayuso, V., Pickles, S., Todd, T., Yue, M., Jansen-West, K., Song, Y., González Bejarano, J., Rawlinson, B., DeTure, M., Graff-Radford, N.R., et al. (2023) TDP-43- regulated cryptic RNAs accumulate in Alzheimer’s disease brains. Mol Neurodegener, 18, 57.

71. Seddighi, S., Qi, Y.A., Brown, A.L., Wilkins, O.G., Bereda, C., Belair, C., Zhang, Y.J., Prudencio, M., Keuss, M.J., Khandeshi, A. et al. (2024) Mis-spliced transcripts generate de novo proteins in TDP-43-related ALS/FTD. Sci Transl Med, 16, eadg7162.

72. Maharana, S., Wang, J., Papadopoulos, D.K., Richter, D., Pozniakovsky, A., Poser, I., Bickle, M., Rizk, S., Guillen-Boixet, J., Franzmann, T.M. et al. (2018) RNA buffers the phase separation behavior of prion-like RNA binding proteins. Science, 360, 918–921.

73. Harini, K., Srivastava, A., Kulandaisamy, A. and Gromiha, M.M. (2022) ProNAB: database for binding affinities of protein-nucleic acid complexes and their mutants. Nucleic Acids Res, 50, D1528–d1534.

74. Ferrari, M.E., Fang, J. and Lohman, T.M. (1997) A mutation in E. coli SSB protein (W54S) alters intra-tetramer negative cooperativity and inter-tetramer positive cooperativity for single-stranded DNA binding. Biophys Chem, 64, 235–251.

75. Zuber, J., Schroeder, S.J., Sun, H., Turner, D.H. and Mathews, D.H. (2022) Nearest neighbor rules for RNA helix folding thermodynamics: improved end effects. Nucleic Acids Res, 50, 5251–5262.

76. Paramanathan, T., Reeves, D., Friedman, L.J., Kondev, J. and Gelles, J. (2014) A general mechanism for competitor-induced dissociation of molecular complexes. Nat Commun, 5, 5207.

77. Velazquez-Campoy, A. and Freire, E. (2006) Isothermal titration calorimetry to determine association constants for high-affinity ligands. Nat Protoc, 1, 186–191.

78. Markova, N. and Hallen, D. (2004) The development of a continuous isothermal titration calorimetric method for equilibrium studies. Anal Biochem, 331, 77–88.

79. Herrera, I. and Winnik, M.A. (2016) Differential Binding Models for Direct and Reverse Isothermal Titration Calorimetry. J Phys Chem B, 120, 2077–2086.

80. Herrera, I. and Winnik, M.A. (2013) Differential binding models for isothermal titration calorimetry: moving beyond the Wiseman isotherm. J Phys Chem B, 117, 8659–8672.

81. Zhao, H., Piszczek, G. and Schuck, P. (2015) SEDPHAT--a platform for global ITC analysis and global multi-method analysis of molecular interactions. Methods, 76, 137–148.

82. Jansson-Fritzberg, L.I., Sousa, C.I., Smallegan, M.J., Song, J.J., Gooding, A.R., Kasinath, V., Rinn, J.L. and Cech, T.R. (2023) DNMT1 inhibition by pUG-fold quadruplex RNA. Rna, 29, 346–360.

83. Baets, J., Duan, X., Wu, Y., Smith, G., Seeley, W.W., Mademan, I., McGrath, N.M., Beadell, N.C., Khoury, J., Botuyan, M.V. et al. (2015) Defects of mutant DNMT1 are linked to a spectrum of neurological disorders. Brain, 138, 845–861.

84. Maresca, A., Del Dotto, V., Capristo, M., Scimonelli, E., Tagliavini, F., Morandi, L., Tropeano, C.V., Caporali, L., Mohamed, S., Roberti, M. et al. (2020) DNMT1 mutations leading to neurodegeneration paradoxically reflect on mitochondrial metabolism. Hum Mol Genet, 29, 1864–1881.

85. Wang, W., Zhao, X., Shao, Y., Duan, X., Wang, Y., Li, J., Li, J., Li, D., Li, X. and Wong, J. (2021) Mutation-induced DNMT1 cleavage drives neurodegenerative disease. Sci Adv, 7, eabe8511.

86. Yang, T., Wei, Q., Pang, D., Cheng, Y., Huang, J., Lin, J., Xiao, Y., Jiang, Q., Wang, S., Li, C. et al. (2024) Clinical and genetic characteristics of ALS patients with variants in genes regulating DNA methylation. J Neurol, 271, 5556–5566.

87. Bayer, C., Pitschelatow, G., Hannemann, N., Linde, J., Reichard, J., Pensold, D. and Zimmer-Bensch, G. (2020) DNA Methyltransferase 1 (DNMT1) Acts on Neurodegeneration by Modulating Proteostasis-Relevant Intracellular Processes. Int J Mol Sci, 21.

88. von Schimmelmann, M., Feinberg, P.A., Sullivan, J.M., Ku, S.M., Badimon, A., Duff, M.K., Wang, Z., Lachmann, A., Dewell, S., Ma’ayan, A., et al. (2016) Polycomb repressive complex 2 (PRC2) silences genes responsible for neurodegeneration. Nat Neurosci, 19, 1321–1330.

89. Wang, X., Goodrich, K.J., Gooding, A.R., Naeem, H., Archer, S., Paucek, R.D., Youmans, D.T., Cech, T.R. and Davidovich, C. (2017) Targeting of Polycomb Repressive Complex 2 to RNA by Short Repeats of Consecutive Guanines. Mol Cell, 65, 1056–1067.e1055.

90. Wang, X., Goodrich, K.J., Conlon, E.G., Gao, J., Erbse, A.H., Manley, J.L. and Cech, T.R. (2019) C9orf72 and triplet repeat disorder RNAs: G-quadruplex formation, binding to PRC2 and implications for disease mechanisms. Rna, 25, 935–947.

91. Song, J., Gooding, A.R., Hemphill, W.O., Love, B.D., Robertson, A., Yao, L., Zon, L.I., North, T.E., Kasinath, V. and Cech, T.R. (2023) Structural basis for inactivation of PRC2 by G-quadruplex RNA. Science, 381, 1331–1337.

92. Appleby-Mallinder, C., Schaber, E., Kirby, J., Shaw, P.J., Cooper-Knock, J., Heath, P.R. and Highley, J.R. (2021) TDP43 proteinopathy is associated with aberrant DNA methylation in human amyotrophic lateral sclerosis. Neuropathol Appl Neurobiol, 47, 61–72.

93. Koike, Y., Sugai, A., Hara, N., Ito, J., Yokoseki, A., Ishihara, T., Yamagishi, T., Tsuboguchi, S., Tada, M., Ikeuchi, T. et al. (2021) Age-related demethylation of the TDP-43 autoregulatory region in the human motor cortex. Commun Biol, 4, 1107.

94. Pacetti, M., De Conti, L., Marasco, L.E., Romano, M., Rashid, M.M., Nubiè, M., Baralle, F.E. and Baralle, M. (2022) Physiological tissue-specific and age-related reduction of mouse TDP-43 levels is regulated by epigenetic modifications. Dis Model Mech, 15.

95. Hou, Y., Dan, X., Babbar, M., Wei, Y., Hasselbalch, S.G., Croteau, D.L. and Bohr, V.A. (2019) Ageing as a risk factor for neurodegenerative disease. Nat Rev Neurol, 15, 565–581.

96. Niccoli, T., Partridge, L. and Isaacs, A.M. (2017) Ageing as a risk factor for ALS/FTD. Hum Mol Genet, 26, R105–R113.

97. Gu, J., Zhang, B., An, R., Qian, W., Han, L., Duan, W., Wang, Z. and Ma, Q. (2022) Molecular Interactions of the Long Noncoding RNA NEAT1 in Cancer. Cancers (Basel), 14.

