## Supplemental Data for "TDP-43 controls RNA structure through high affinity lattice interactions"

**Supplementary Data**

Rahul Vivek,[1] Takuma Kume,[1] Saeed Roschdi,[1] M. Thomas Record, Jr.,[2, 3] Aaron A. Hoskins, [1,3] and Samuel E. Butcher [1*]

[1] Department of Biochemistry, University of Wisconsin-Madison, Madison, WI, USA - 53706.

[2] Department of Biochemical Sciences, University of Wisconsin-Madison, Madison, WI, USA – 53706.

[3] Department of Chemistry, University of Wisconsin-Madison, Madison, WI, USA – 53706.

**Supplementary Tables:**

Supplementary Table 1: TDP-43 Binding Constants for $(GU)_6$

| TDP-43 Domains | Incubation Time (h) | Apparent $K_D$ [pM] |
|---|---|---|
| RBD | 0.5 | 72 ± 5 |
| | 2 | 63 ± 6 |
| | 6 | 70 ± 5 |
| | 12 | 73 ± 5 |
| NTD-RBD | 0.5 | 266 ± 18 |
| | 2 | 245 ± 30 |
| | 6 | 237 ± 23 |
| | 12 | 251 ± 17 |

Supplementary Table 2: Kinetic analysis of TDP-43 RBD and NTD-RBD dissociation and exchange on $(GU)_6$

| Domains | $k_{-1}$ ($x\ 10^{-2}$) ($s^{-1}$) | $k_{exch}$ ($x10^3$) ($M^{-1}\ s^{-1}$) | *Apparent* $K_D$ [pM] | Estimated $k_1$ (x $10^8$) $M^{-1}\ s^{-1}$ |
|---|---|---|---|---|
| RBD | 3.0 ± 2.1 | 4.9 ± 1.7 | 73 ± 5 | 4.1 ± 2.9 |
| NTD-RBD | 1.6 ± 0.3 | 9.0 ± 0.4 | 251 ± 17 | 0.6 ± 0.1 |

Supplementary Table 3: RNA-binding proteins from *Homo sapiens* with high-affinity interactions

| Protein | RNA | Method | $K_D$ [pM] |
|---|---|---|---|
| VEGF165 | AF83-7 | Bio-layer interferometry | 1 |
| VEGF165 | AF83-19 | Bio-layer interferometry | 1 |
| Prothrombin | AF113-18 | Bio-layer interferometry | 1.8 |
| Antiviral innate immune response receptor RIG-I | 5'ppp 14-bp-dsRNA-F | ATPase coupled binding assay | 20 |
| U1 small nuclear ribonucleoprotein A | Stem-loop 2 RNA | EMSA | 30 |
| U1 small nuclear ribonucleoprotein A | U1 hairpin II | Surface plasmon resonance | 32 |
| Protein lin-28 homolog A | let-7 | EMSA | 33 |
| U1 small nuclear ribonucleoprotein A | U1 hairpin II | Surface plasmon resonance | 34 |
| U1 small nuclear ribonucleoprotein A | U1 hairpin II | Surface plasmon resonance | 40 |
| U1 small nuclear ribonucleoprotein A | U1 hairpin II | EMSA | 47 |
| U1 small nuclear ribonucleoprotein A | stem-loop II (SLII) RNA | Filter binding | 50 |
| Prothrombin | Pig-10 RNA aptamer | Filter binding | 50 |
| Antiviral innate immune response receptor RIG-I | 14-bp-dsRNA-F | Fluorescence anisotropy | 50 |
| RNA binding protein fox-1 homolog 1 | RNA | Surface plasmon resonance | 60 |
| Pumilio homolog 1 | NRE- containing RNA | EMSA | 60 |
| U1 small nuclear ribonucleoprotein A | U1 hairpin II | Surface plasmon resonance | 70 |
| PUM2 | hbNRE2 | EMSA | 70 |
| Pumilio homolog 2 | p38aNREb | EMSA | 80 |
| Helicase-RD | Cap-0 HP RNA | ATPase coupled binding assay | 80 |
| Prothrombin | Toggle-25 RNA aptamer | Filter binding | 83 |

Search Query for Binding Affinity Range: 1 fM to 100 pM ($K_D$)
Extracted from the proNAB database

**Supplementary Figure 1**

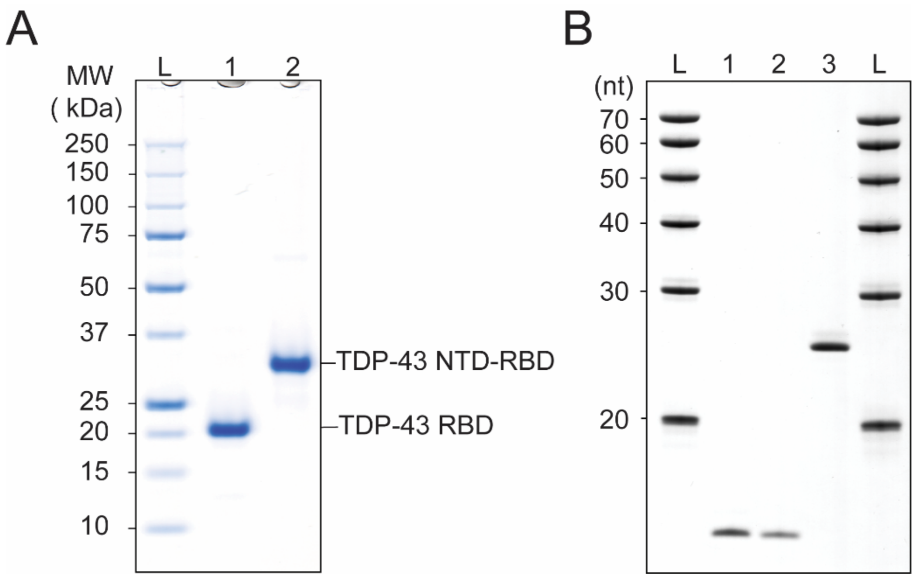


**Supplementary Figure 1. Purity of TDP-43 protein and RNA used in the study.**

(A) SDS-PAGE analysis of purified TDP-43 RBD and NTD-RBD on NuPAGE™ 4-12% Bis-Tris Mini Protein Gels (1.0 mm). Precision Plus Protein™ Unstained Standards were used as molecular weight markers. Lanes: L, protein ladder; 1, TDP-43 RBD (19.9 kDa); 2, TDP-43 NTD-RBD (30.8 kDa). Each protein lane contained 0.5 µg of protein. The gel was stained with Coomassie Brilliant Blue. (B) Denaturing 20% PAGE analysis of purified RNAs. Lanes: L, RNA ladder; 1, AUG12 (12 nt); 2, $(GU)_6$ (12 nt); 3, $(GU)_{12}$ (24 nt). Each lane contained 10 µL of RNA at 15 µM, except for $(GU)_6$, which was loaded at 10 µM. The gel was stained with toluidine blue.

**Supplementary Figure 2**

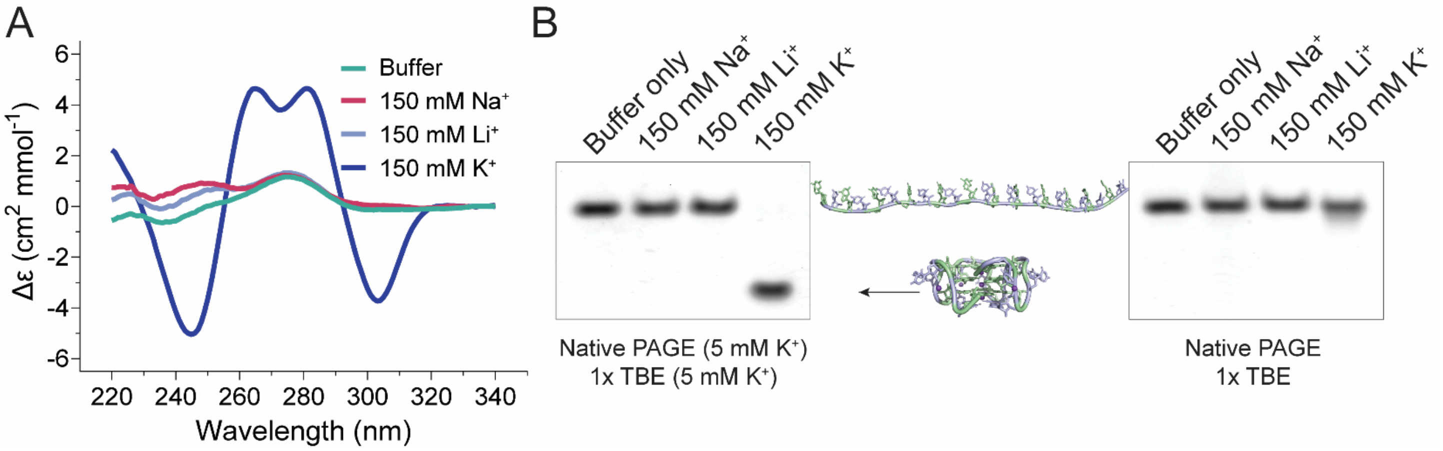


**Supplementary Figure 2. pUG RNA folds only in the presence of potassium ions.**

(A) CD spectra of 20 µM $(GU)_{12}$ RNA in 20 mM Bis-Tris (pH 7.0) containing no ions (green), or 150 mM $Na^+$ (red), 150 mM $Li^+$ (light blue), or 150 mM $K^+$ (dark blue). (B) Native 8% polyacrylamide gel showing the mobility of 20 µM $(GU)_{12}$ RNA in buffers containing different ions. Gels were run either in the presence (left) or absence (right) of 5 mM $K^+$ in both the gel and running buffer. RNA was visualized by toluidine blue staining.

**Supplementary Figure 3**

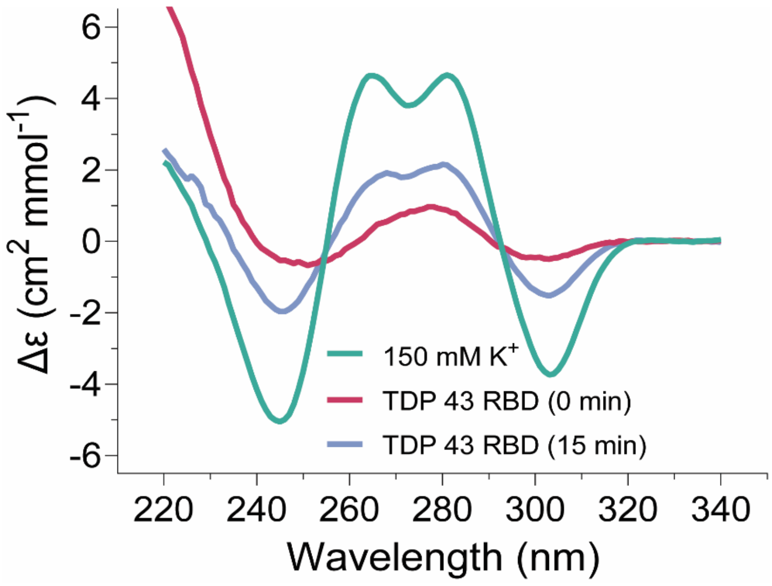


**Supplementary Figure 3: CD spectra of $(GU)_{12}$ post kinetic experiments.**

CD spectra of $(GU)_{12}$ RNA 3 h after kinetic measurements. Cyan, RNA + 150 mM $K^+$ injected at time = 0 min. Red, RNA after simultaneous injection of $K^+$ ions and TDP-43 RBD at time = 0 min. Blue, $K^+$ injection at time = 0 min. and TDP-43 RBD injection at time = 15 min. Protein-only signals were measured and subtracted from the raw CD data to isolate the RNA spectra.

**Supplementary Figure 4**

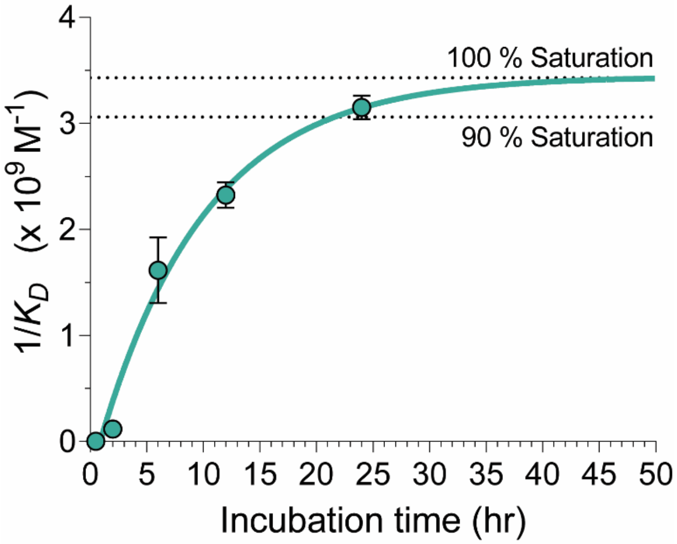


**Supplementary Figure 4. Time-dependent equilibration of TDP-43 RBD binding to $(GU)_{12}$ RNA.**
Plot shows the change in apparent binding affinity ($1/K_D$) of TDP-43 RBD for $(GU)_{12}$ RNA as a function of incubation time. The apparent affinity increases over time, reaching ~94% of equilibrium after 24 h and approaching 99% equilibrium by ~43 h. Data were fit using a single-exponential nonlinear regression model, yielding a $K_D$ of 290 ± 41 pM at complete saturation.

**Supplementary Figure 5**

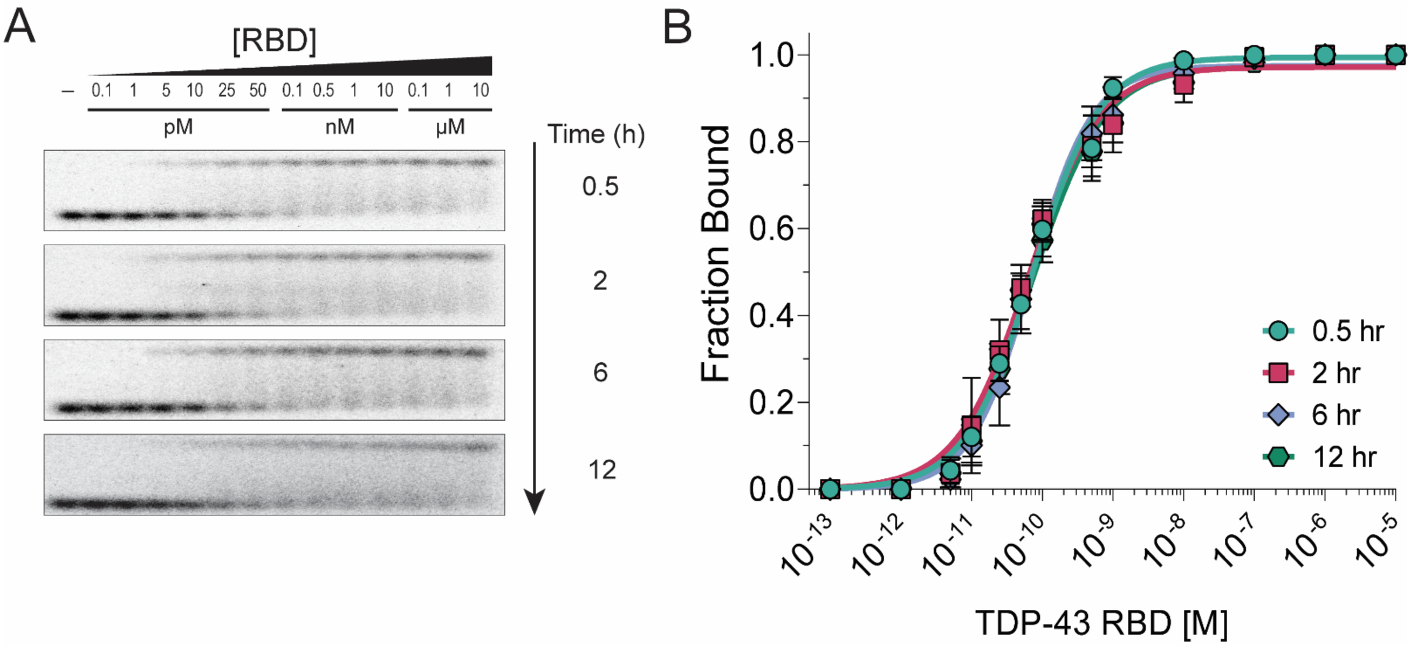


**Supplementary Figure 5. Quantitative analysis of TDP-43 RBD binding to $(GU)_6$ RNA**

(A) EMSA showing complex formation between TDP-43 RBD and $(GU)_6$ RNA at different incubation times prior to electrophoresis. (B) Binding curves derived from quantification of EMSA data at indicated incubation times.

**Supplementary Figure 6**

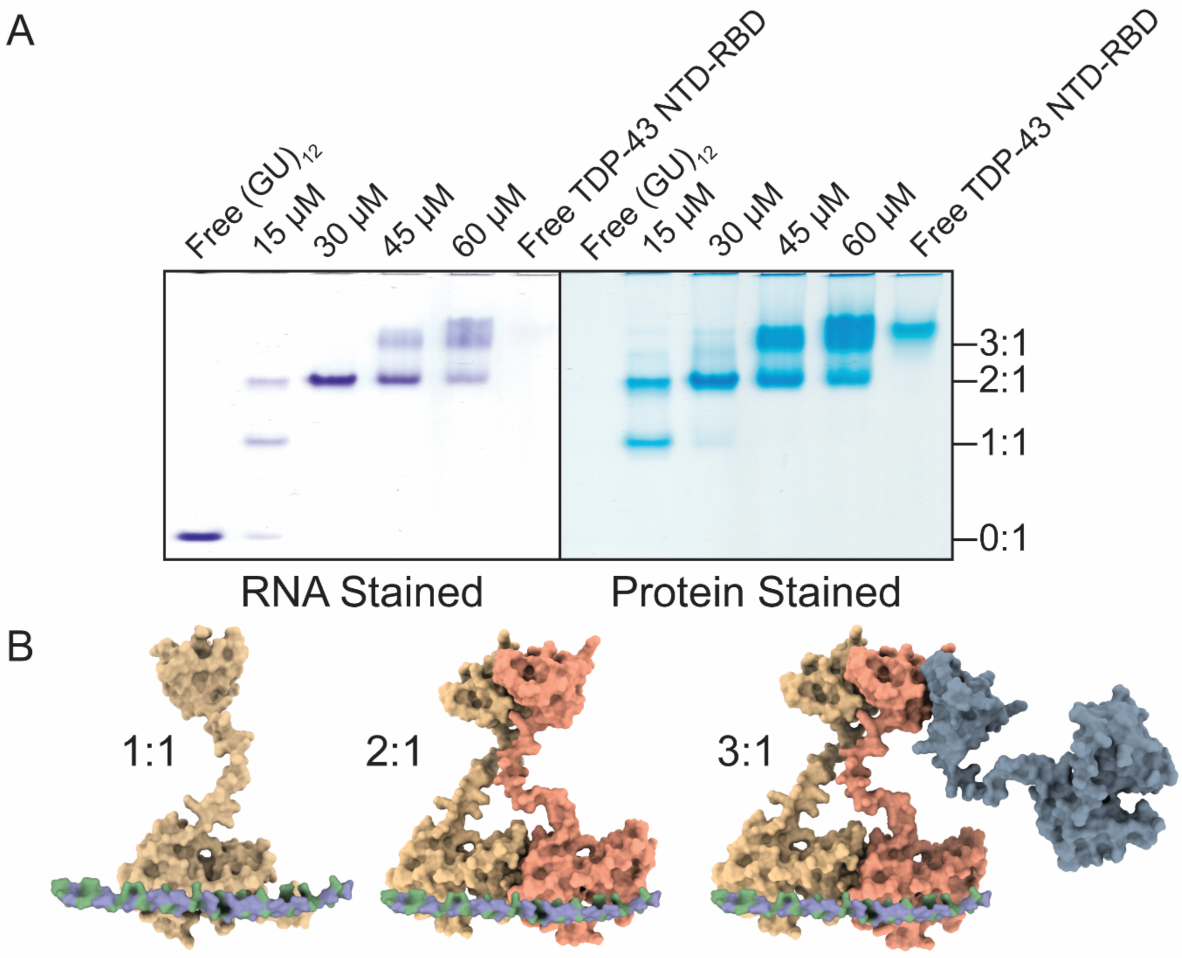


**Supplementary Figure 6. Stoichiometric binding of TDP-43 NTD-RBD to $(GU)_{12}$ RNA.**

A. Native 8% polyacrylamide gels showing complexes formed between 15 µM $(GU)_{12}$ RNA and TDP-43 NTD-RBD. The left panel is RNA-stained with toluidine blue, and the right panel is protein-stained with Coomassie Brilliant Blue (CBB). Binding buffer: 20 mM Bis-Tris (pH 7.0), 150 mM $Li^{+}$, 20% sucrose. The Free TDP-43 NTD-RBD lane contains 15 µM Protein. B. AlphaFold models for the annotated complexes (1:1, 2:1, and 3:1 protein–RNA complexes).

**Supplementary Figure 7**

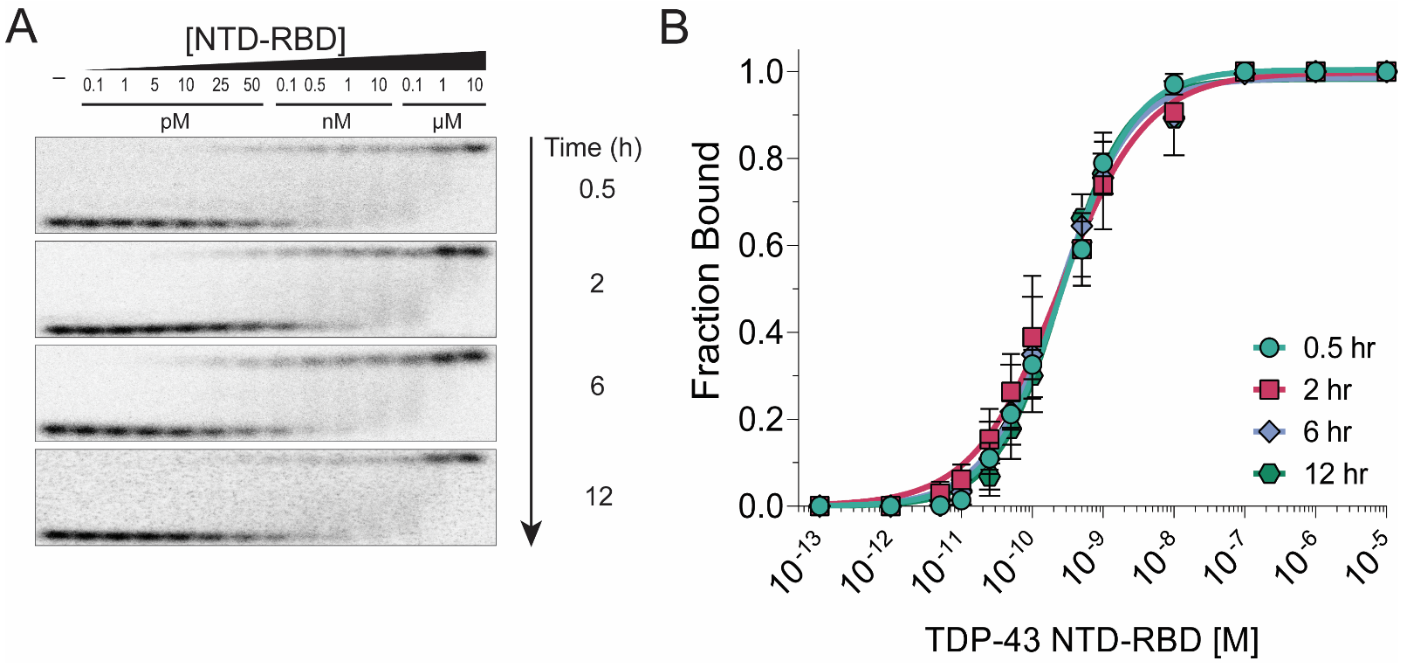


**Supplementary Figure 7. Quantitative analysis of TDP-43 NTD-RBD binding to $(GU)_6$ RNA**

(A) EMSA showing complex formation between TDP-43 NTD-RBD and $(GU)_6$ RNA at different incubation times prior to electrophoresis. (B) Binding curves derived from quantification of EMSA data at indicated incubation times.

**Supplementary Figure 8**

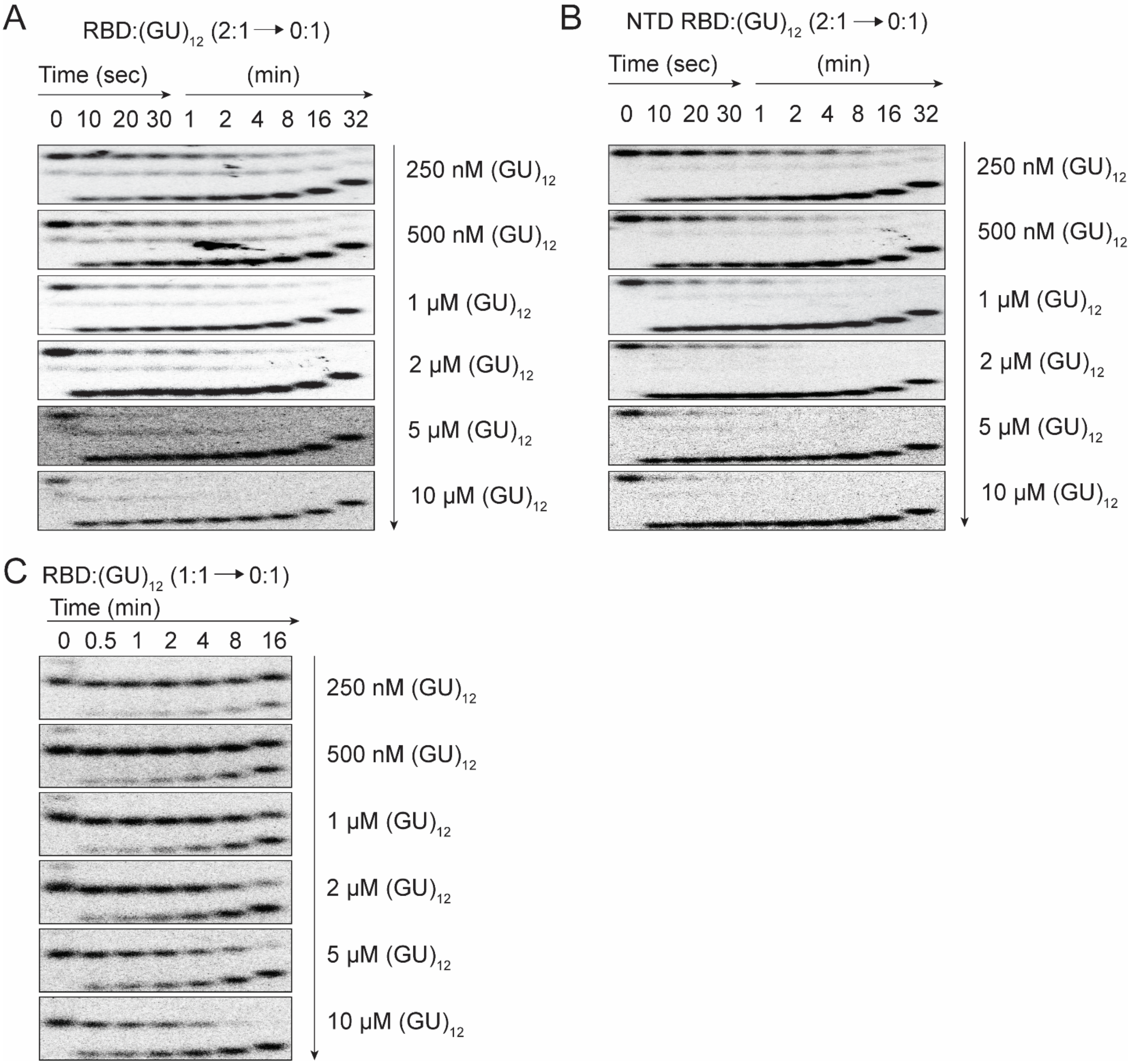


**Supplementary Figure 8. Dissociation kinetics of TDP-43 RBD and NTD-RBD with $(GU)_{12}$ RNA.**
(A-C) EMSAs showing dissociation of pre-formed 10 nM TDP-43–RNA complexes using 10 pM $^{32}P$-labeled $(GU)_{12}$ RNA. (A) 2:1 RBD, (B) 2:1 NTD-RBD and (C) 1:1 RBD. The 2:1 complexes were prepared by incubating protein and RNA for 12-16 h, whereas the 1:1 RBD complex was obtained by incubation for 0.5 h. Complexes were challenged with increasing concentrations of unlabeled $(GU)_{12}$ RNA (250 nM - 10 μM).

**Supplementary Figure 9**

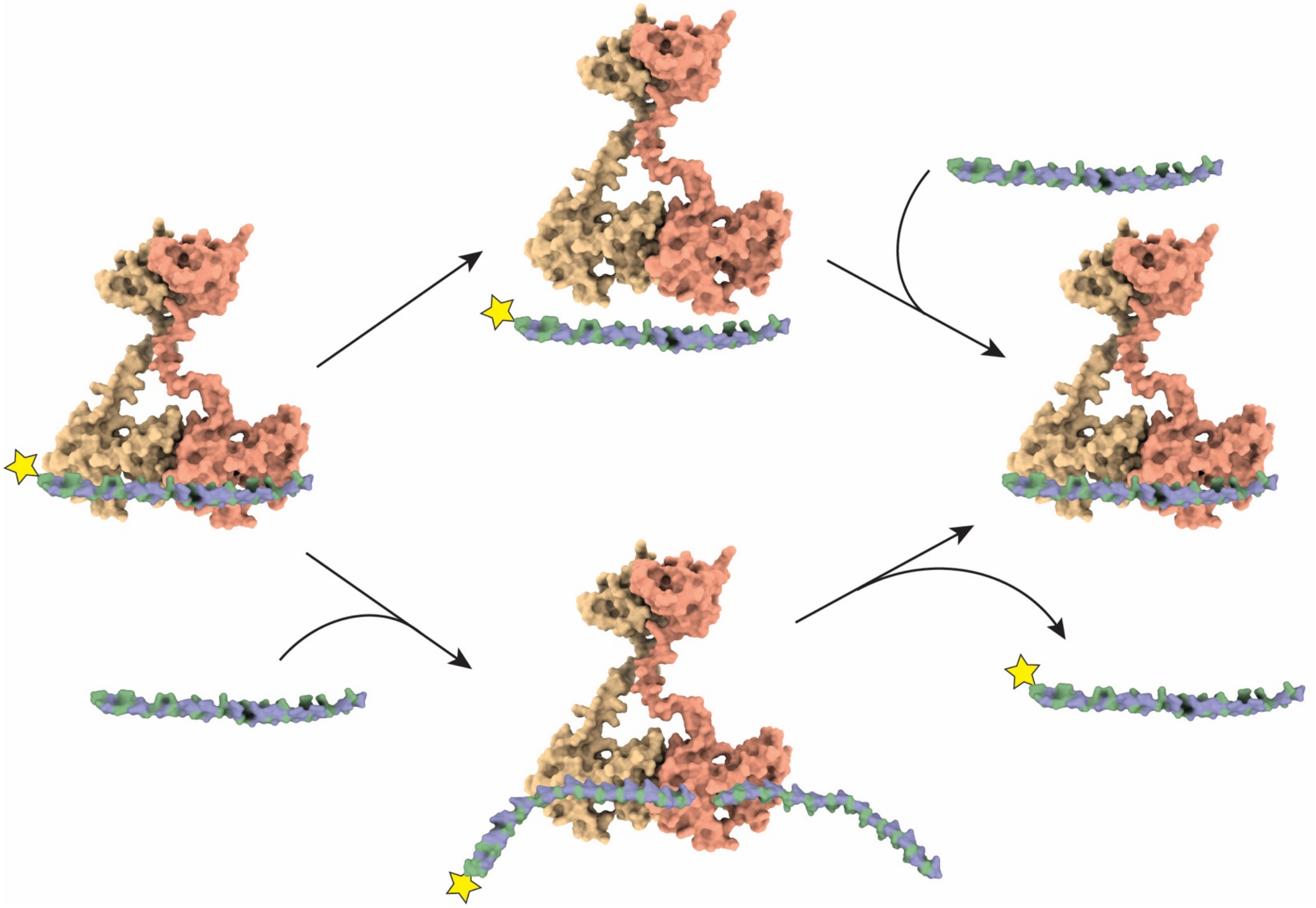

**Supplementary Figure 9: Two possible pathways for facilitated dissociation of $(GU)_{12}$.**

AlphaFold models illustrating the exchange of $(GU)_{12}$ RNA at higher competitor concentrations for NTD-RBD. A similar process occurs for RBD (not shown). The yellow star denotes $^{32}P$ labeled $(GU)_{12}$. Top pathway corresponds to exchange facilitated by micro-dissociation, lower pathway corresponds to direct transfer involving a ternary complex.

**Supplementary Figure 10**

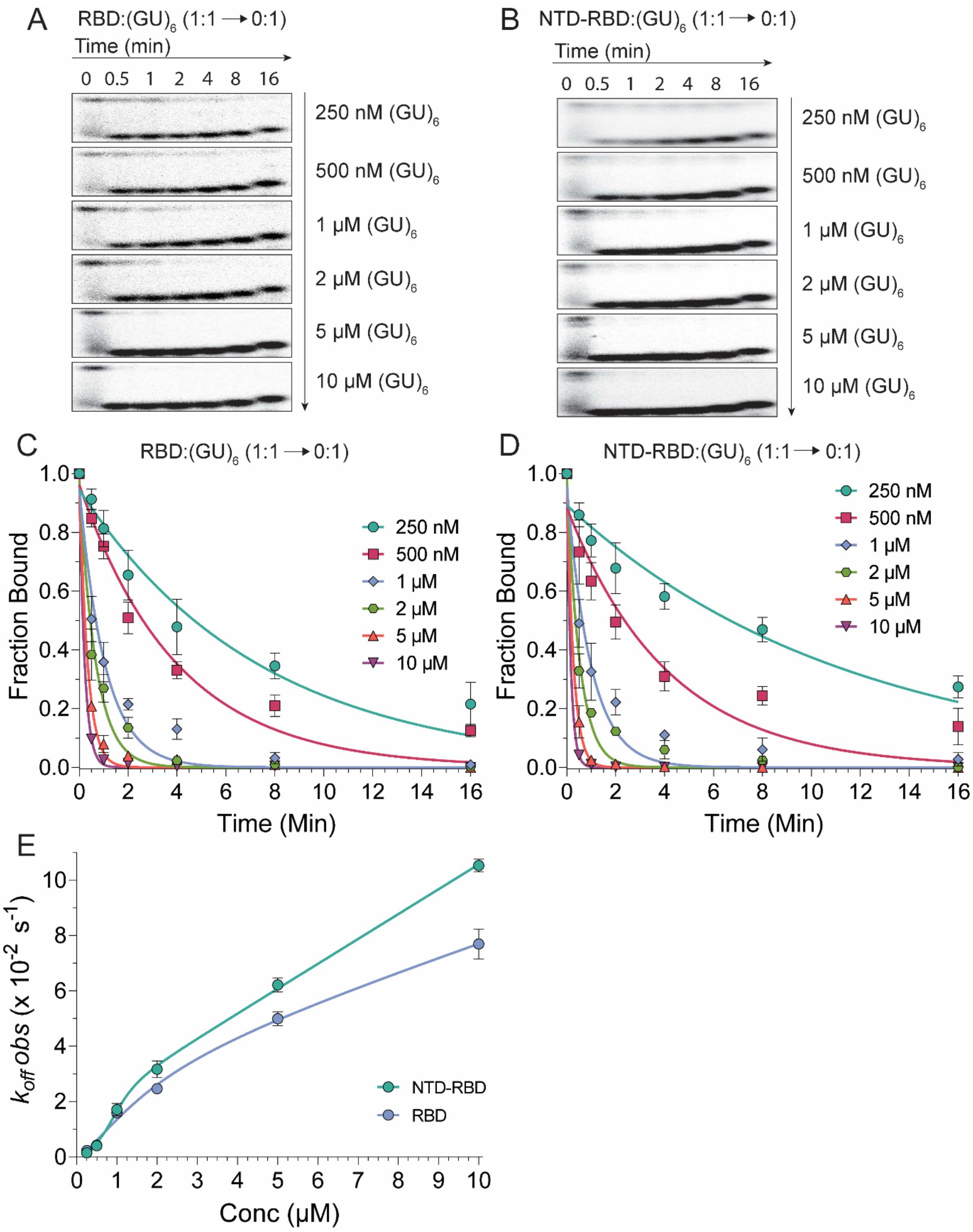


**Supplementary Figure 10. Dissociation kinetics of TDP-43 RBD and NTD-RBD complexes with $(GU)_6$ RNA.** (A, B) EMSAs showing dissociation of complexes formed between 10 pM $^{32}$P-labeled $(GU)_6$ RNA and either 10 nM TDP-43 RBD (A) or 10 nM NTD-RBD (B). Complexes were challenged with increasing concentrations of unlabeled $(GU)_6$ RNA (250 nM - 10 μM). (C, D) Dissociation kinetics for RBD (C) and NTD-RBD (D). (E) Apparent

$k_{obs}$ values plotted against competitor RNA concentration and fitted with a biphasic model for classical and facilitated dissociation.

**Supplementary Figure 11**

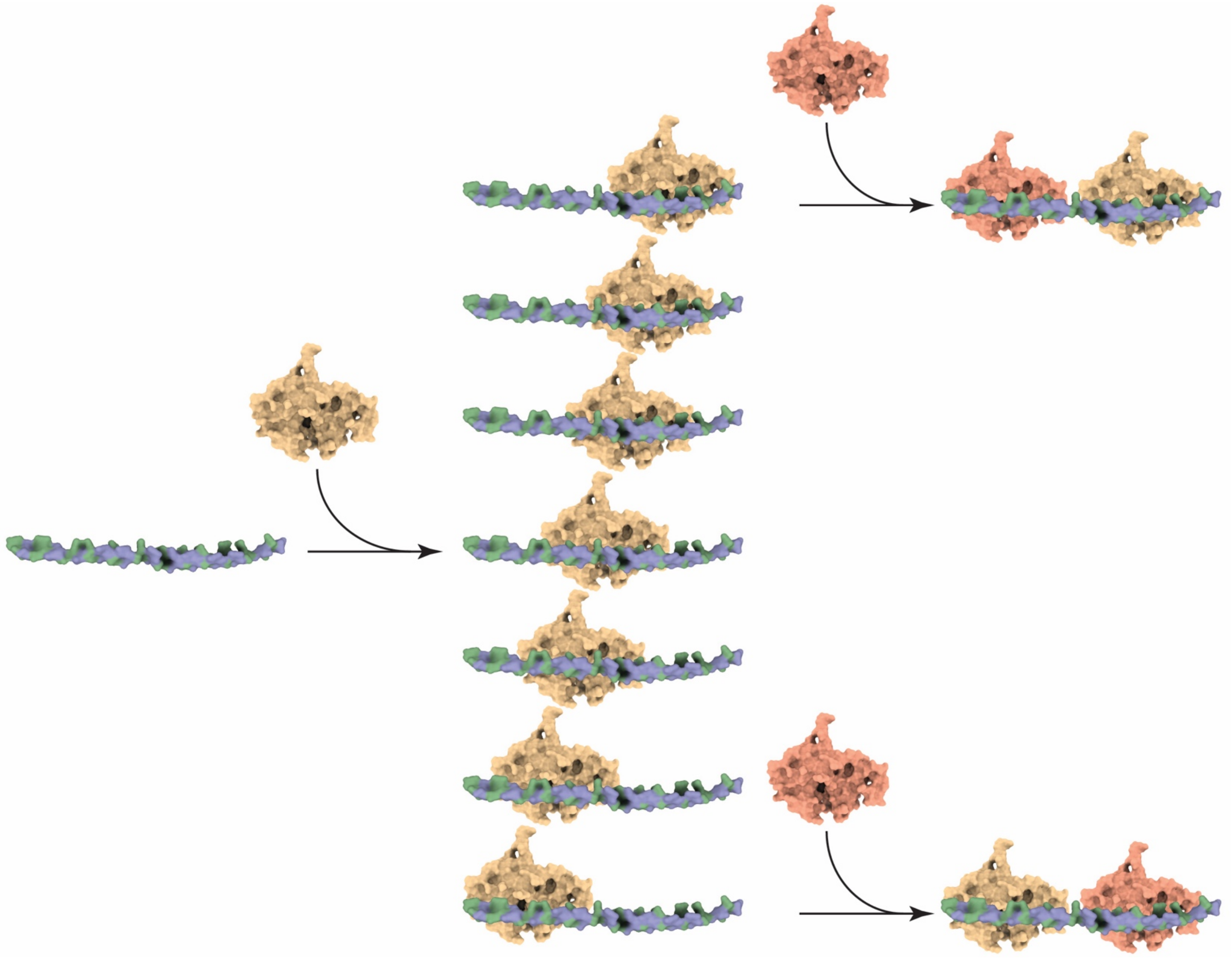

**Supplementary Figure 11. Longer pUG RNAs present multiple overlapping TDP-43 binding sites.**
The 24-nt $(GU)_{12}$ RNA contains seven overlapping binding sites, which increase the fraction bound for the first event by raising the effective concentration of available sites at equilibrium. 5 out of 7 initial binding events exclude binding of a second molecule.
